# Chemokine-Dependent Natural Killer Cells Prevent Pulmonary Metastasis in a Newly Established Mouse Osteosarcoma Model

**DOI:** 10.64898/2026.07.19.738728

**Authors:** Jifeng Yang, Frederique Zindy, Shaela Fields, Judith Hyle, Laura J. Janke, Qianqian Li, Beisi Xu, Qiong Zhang, Gang Wu, Ti-Cheng Chang, Monika Wierdl, Lillian M. Guenther, Linda M. Hendershot, Martine F. Roussel, Charles J. Sherr, Chunliang Li

## Abstract

Rampant genomic instability of osteosarcoma (OS) and associated inter- and intratumoral heterogeneity convey a high risk of metastasis despite contemporary multimodal therapy. A mouse primary OS tumor model, originating from Myc-overexpressing *Trp53*-null mesenchymal progenitor cells, closely mimics cardinal genetic features and gene expression patterns of human OS samples. Subcutaneous injection of OS cells into immunocompromised NSG mice produced extensive metastases, whereas many fewer metastases occurred in T cell-deficient nude mice, suggesting a principal role for innate immunity in controlling OS dissemination. Depletion of natural killer (NK) cells in nude mice facilitated OS metastasis. OS cells released a suite of chemokines, with CCL2 the most prominent. A genome-wide CRISPR/Cas9 screen in OS cells identified 11 genes, including *Ccl2*, whose loss facilitated pulmonary metastasis in nude mice. Disruption of CCL2 in OS cells partially phenocopied the effects of antibody-dependent NK cell depletion, underscoring a plausible OS-NK signaling pathway that limits OS metastasis.

**Significance:** Osteosarcoma exhibits complex genomic instability and a propensity for pulmonary metastasis that limit chemotherapeutic response and patient survival. Despite the plethora of heterogeneous genetic alterations that connote poor prognosis, a novel preclinical *in vivo* model for studying metastasis highlights potentially targetable signaling between osteosarcoma cell-derived chemokines and pulmonary NK cells.

## Introduction

Osteosarcoma (OS), the most common primary bone malignancy in children and young adults, typically arises in the metaphysis of growing long bones, most frequently the humerus and femur (1). An estimated 1,000 new cases are diagnosed each year in the United States. The five-year survival of OS patients after tumor resection alone is less than 20%, limited by the presumed expansion of pre-surgical tumor micro-metastases primarily to the lung. The addition of systemic chemotherapy raised the five-year survival rate for patients with localized OS to 60-70%, but without further improvement for more than four decades. Metastatic and relapsed disease remains the primary cause of death in one-third of treated patients (2, 3). A major feature of OS is genomic instability and inter- and intratumoral heterogeneity (4) due to ongoing chromothripsis in at least 75% of primary tumors (5, 6), which leads to challenges in developing targeted therapies for recurrent lesions and drives metastatic clonal selection (7–9). Moreover, the tumor microenvironment (TME) of OS is highly immunosuppressive, and despite widespread interest in potential immunotherapies, none have been approved to date for OS treatment (10). Although treatments targeting candidate driver genes can reduce bulk tumor burden, there are no as-yet unifying mechanisms that clearly explain metastatic spread in OS (8).

Basic and preclinical studies have generally relied on a small subset of established OS cell lines, which were adapted to *in vitro* culture conditions and continuous passage, with only a few consistently forming metastases when injected orthotopically or intravenously into mice (11, 12). Among six routinely studied OS cell lines arising in inbred (Balb/c, C3H, and C57BL/6J) mice, only half metastasized to the lung following intravenous or orthotopic transplantation into syngeneic hosts (13), with similar variability observed with patient-derived xenografts (PDXs) in immune-compromised <u>N</u>OD-<u>s</u>cid IL2R <u>g</u>amma chain-null (NSG) mice (14). Notably, intravenous injection of those OS lines that can generate lung metastases fails to recapitulate the biological processes of local dissemination, intravasation, and colonization that characterize the metastatic spread of primary tumors. Moreover, transcriptomic and epigenetic profiling revealed substantial differences among human OS models, with copy-number changes and structural variants predominating over relatively few point mutations in protein-coding genes (8). Although PDXs can recapitulate the genotypes of the tumors from which they arise (14), the genomic heterogeneity and selection have limited mechanistic studies of metastasis in particular. Moreover, transplantation models of PDXs generally rely on immunodeficient NSG host mice, which lack both adaptive and innate immune cells. Differences in metastatic potential likely reflect not only intrinsic variation among OS samples but also diverse extrinsic TMEs across sites and species (15, 16). Genetically engineered mouse models that fully recapitulate human OS generally exhibit a relatively long time to onset and metastasis (17–19), implying ongoing clonal selection.

Despite the complexity and variability of OS genomes, high-throughput genomic studies have revealed that *TP53* loss-of-function (around 70% of cases) and *MYC* gene amplification (25-50% across studies) are the most prevalent mutations in human OS (20, 21). Here, we report the establishment of a reproducible mouse OS model using the *Myc/Trp53* gene combination, which fully recapitulates human OS pathogenesis and transcriptomic signatures. Cell lines derived from primary OS tumors can be continuously cultured, are metastatic, and despite their genetic complexity and near-tetraploid chromosomal content, are amenable to genome editing. Leveraging this new OS model through experimental immune cell perturbation and functional genomics revealed that NK cells and their crosstalk with chemokine-secreting tumors are particularly critical for controlling OS pulmonary metastasis.

## Results

### A Myc-driven, *Trp53*-null OS mouse model

Granular neural progenitors (GNPs) dispersed from the developing cerebella of postnatal day-7 (P7) *Trp53*-null C57BL/6J mice and enriched by Percoll gradient centrifugation, were engineered to robustly express *Myc;* these cells reproducibly form medulloblastomas (MBs) when injected into the cerebral cortices of CD-1 nude mice (22). Combined disruption of *Trp53* and *Myc* overexpression was required for tumorigenesis, with no tumors arising from *Trp53*-null or *Myc*-overexpressing GNPs alone. Although injection of these tumor-initiating cells deep into the cerebral cortices of recipient mice invariably produced MBs, we subsequently observed that cells inadvertently injected just below the calvarium generated bone tumors (23, 24). Moreover, when the Percoll-purified, GNP-enriched fraction was transiently expanded in culture to form presumptive neurospheres, proximal subcalvarial injection of pooled *Myc* retrovirus-infected, *Trp53*-null sphere cells into the brains of CD-1 nude recipients stably produced OS (Schematic **Fig. 1A**). In unrelated experiments, we engineered a carboxy-terminally tagged *Myc* allele encoding a mini auxin-inducible degron (miniAID)(25) that phenocopied the rapid turnover rate (data not shown) and oncogenic function of the wild-type (WT) MYC protein in the absence of auxin treatment. Both the overexpressed WT and miniAID-tagged *Myc* alleles collaborated with *Trp53* loss to form bone tumors (25/33, ∼76%) or MB (6/33, ∼18%)(**Fig. 1B and 1C; Supplementary Table S1**). These cartilage-containing and mineralized bone tumors (**Fig. 1D-1G**) robustly expressed the osteoblast marker SATB2 (**Fig. 1H**), and 36% (9 of 25) of primary bone tumors showed low or loss of PTEN expression (**Fig. 1I and 1J; Supplementary Table S1**). Five randomly selected individual primary bone tumors were isolated and stably cultured *in vitro*.

**Fig. 1.**
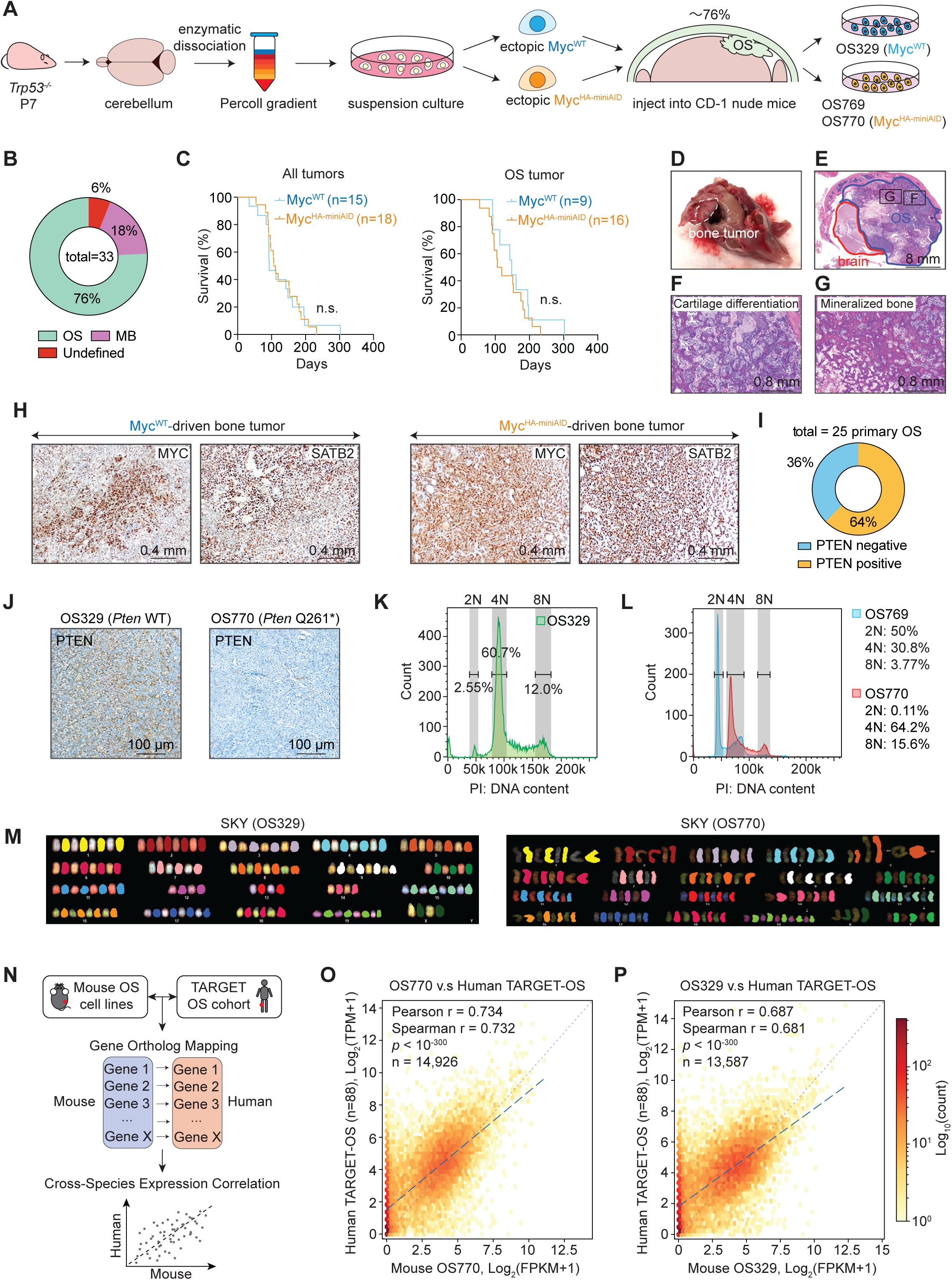
A newly established Myc-driven OS model recapitulates human OS signatures. (A) Schematic diagram of deriving a primary mouse Myc-driven OS model. Cells from the postnatal day 7 (P7) cerebellums of *Trp53*-null C57BL/6J mice were enzymatically dissociated. The cell mixture was then fractionated by the Percoll gradient purification to enrich GNPs. Enriched GNPs cultured in suspension (sphere cells) were infected with retroviral vectors encoding wild-type (WT) Myc (Myc^WT^) or HA-miniAID-tag Myc (Myc^HA-miniAID^). Cells with high ectopic Myc expression were sorted and injected into the cerebral cortices of CD-1 nude host mice. Cells injected beneath the calvaria formed OS tumors that were stably cultured *in vitro*. (B) Pie chart summary for the percentages of all tumor types generated by the above-mentioned method. (C) Overall survival curve of tumor-bearing CD-1 nude mice injected with sphere cells carrying either ectopic Myc^WT^ (blue) or Myc^HA-miniAID^ (orange). Kaplan-Meier survival curves for all tumors (including MBs) and OS only are plotted separately. Statistical analysis was performed using a log-rank test. Non-significant (n.s.) *p* > 0.05. (D) A typical brain image of a CD-1 nude recipient mouse containing a bony tumor. (E) Hematoxylin and Eosin (H&E) staining of a typical primary bone tumor section (outlined in blue) alongside brain tissues (outlined in red). Boxes represent areas shown in (F) and (G). (F) Image of H&E-stained primary bone tumor containing classic differentiated cartilage. (G) Image of H&E-stained primary bone tumor containing classic mineralized bone. (H) Primary OS tumor cells expressing MYC and SATB2 were detected by immunohistochemistry. (I) Pie chart showing the percentages of OS primary tumors with PTEN status defined by immunohistochemistry staining against PTEN protein. (J) Representative immunohistochemical images of primary OS tumor cells defined as PTEN-positive and PTEN-negative based on PTEN protein expression. (K) DNA content of OS329 cells was quantified by PI staining and flow cytometry. (L) DNA content of OS769 and OS770 cells was quantified by PI staining and flow cytometry. (M) Fluorescence-labeled spectral karyotyping (SKY) analysis of OS tumor cell lines OS329 and OS770. Chromosomes recognized by different probes are displayed in distinct colors. (N) Schematic diagram of cross-species transcriptome correlation analysis. RNA-seq expression data from OS cells were compared against bulk RNA-seq from 88 pediatric OS patients in the TARGET-OS cohort. Expression values were log_2_(TPM+1)-transformed independently for each species. One-to-one orthologous gene pairs were retrieved from Ensembl BioMart (release 113, GRCm39/GRCh38; ortholog_one2one, confidence = 1), yielding 14,926 pairs between OS770 and human OS and 13,587 pairs between OS329 and human OS after intersection. Pearson correlation, Spearman correlation, and linear regression lines were calculated from mean expression values per gene. (O) The OS770 cell line showed a relatively high transcriptional correlation with human OS samples (Pearson r = 0.734; Spearman r = 0.732; *p* < 10^-300^). (P) The OS329 cell line showed a relatively high transcriptional correlation with human OS samples (Pearson r = 0.687; Spearman r = 0.681; *p* < 10^-300^).

Although *Trp53*-null, *Myc*-overexpressing tumor-initiating sphere cells were diploid, two of the recovered OS tumors (OS770 and OS329) that returned to culture were near-tetraploid, while another one (OS769) retained a diploid karyotype (**Fig. 1K-1M**). Analysis of 20 metaphases from each of the two near-tetraploid cell lines showed 69-78 chromosomes per metaphase. The chromosomes of both OS770 and OS329 cells exhibited distinct subclonal populations, each containing chromosomal deletions, duplications, and translocations (**Supplementary Fig. S1A-B**). Whole-genome sequencing (WGS) and RNA sequencing (RNA-seq) of OS770 further revealed the acquisition of a duplicated (four copies) nonsense mutant *Pten* (Q261*)(**Supplementary Fig. S1C and S1D**) and *Myc* retroviral integrations across different chromosomes (**Supplementary Table S2**), suggesting strong selection for both events during tumor initiation and cell line establishment. Cross-species transcriptomic comparison revealed that the representative OS770 and OS329 cells (**Fig. 1N-1P**) closely recapitulated the transcriptional signatures of clinical OS patient samples (from the Therapeutically Applicable Research to Generate Effective Treatments, TARGET cohort)(26).

Regarding the presumptive tumor cells of origin, we inferred that, in addition to the majority of authentic GNPs that formed MBs, the cell fraction enriched by Percoll gradient centrifugation also contained a small fraction of *Trp53*-null, *Myc*-overexpressing mesenchymal progenitor cells capable of forming OS (**Supplementary Fig. S2A**). Single-cell RNA-seq analysis of 4,707 cells from the freshly GNP-enriched fraction identified several distinct cell populations, including 60.4% GNPs, 18.7% granule neurons, 9.6% interneuron progenitors, 9.5% GABAergic interneurons, and small populations of Bergmann glia and microglia cells. Culturing *Trp53*-null/*Myc*-overexpressing GNP-enriched fraction in a cytokine-supplemented (EGF and FGF) sphere suspension system *in vitro* led to the depletion of (neuronally marked) *Atoh1-*positive GNPs with a concordant expansion of a minor *Atoh1*-negative population. Moreover, the freshly GNP-enriched fraction contained a *Pdgfra*-positive (mesenchymal progenitor marker) cell population, a subset of which was retained in sphere cultures. Ectopic MYC expression further reprogrammed and expanded the sphere cells, leading to the emergence of an *Atoh1*-negative, *Satb2/Alpl*-positive (osteoblast markers) population (**Supplementary Fig. S2B and S2C)**.

Lineage tracing was further conducted by crossing an *Atoh1-Cre^ERT2^* mouse strain with the *Rosa-LoxP-td<u>T</u>omato-stop codon-LoxP-<u>G</u>FP* (*mT/mG*) transgenic strain in a *Trp53*-null background to obtain animals expressing both fluorescent reporter alleles. After *in vivo* tamoxifen treatment, tumor-initiating cells were again recovered and returned to sphere culture. Under these conditions, *Atoh1-*positive cells excised the tdTomato cassette and expressed GFP, whereas *Atoh1-*negative cells remained tdTomato^+^/GFP^-^ (**Supplementary Fig. S2D**). Flow cytometric analysis confirmed the rapid depletion of the *Atoh1*-positive neuronal lineage in sphere cultures (**Supplementary Fig. S2E**), accompanied by the preferential expansion of *Atoh1*-negative cells of OS origin arising from a distinct cellular population. In summary, we inadvertently generated a mouse OS model by successfully targeting engineered mesenchymal progenitor cells to a subcalvarial bone niche.

### Characterization of OS metastatic potential

In patients with OS, metastases occur primarily in the lungs and at distal bony sites, though more advanced and end-stage patients may develop overt metastases in other organs. Subcutaneous injections of 3 × 10^6^ OS770 cells into NSG mice, which are deficient in both adaptive and innate immunity, induced flank tumors that reached the humane endpoint for sacrifice after 3-4 weeks post-injection due to extensive tumor burdens. When necropsied, all NSG host mice demonstrated extensive lung metastatic nodules expressing MYC and SATB2 proteins (**Fig. 2A-2C; Supplementary Fig. S3A and S3B**). Extensive scans of histopathological sections from each animal further identified widespread micro-metastases in the livers, axial lymph nodes, adrenal glands, and kidneys of NSG hosts (**Supplementary Fig. S3C-S3F**). Similar metastatic capacity was observed when OS329 cells were injected subcutaneously into NSG hosts (**Supplementary Fig. S4A-S4F**). Both OS770 and OS329 cells maintained a near-tetraploid karyotype from the time of subcutaneous engraftment through lung metastatic colonization in NSG hosts, indicating that the near-tetraploid state is preserved throughout *in vivo* selection (**Supplementary Fig. S5A-S5E**). Remarkably, when OS cells were subcutaneously injected into immunocompetent syngeneic C57BL/6J mice or T cell-deficient nude mice, no visible lung metastases were observed, despite rapid flank tumor development. Histopathological review identified a few micro-metastases in the lungs of some nude mice, and none in C57BL/6J mice (**Fig. 2A-2C; Supplementary Fig. S4A**). Consistently, primary subcutaneous tumors grew rapidly in the flanks of both nude and NSG mice, necessitating their sacrifice 2-4 weeks post-injection (**Fig. 2D**). Bioluminescent imaging demonstrated that robust lung metastatic signals were detected as early as 13 days post-subcutaneous injection of luciferase-labeled OS770 cells in NSG mice, but not in CD-1 nude mice (**Fig. 2E and 2F**).

**Fig. 2.**
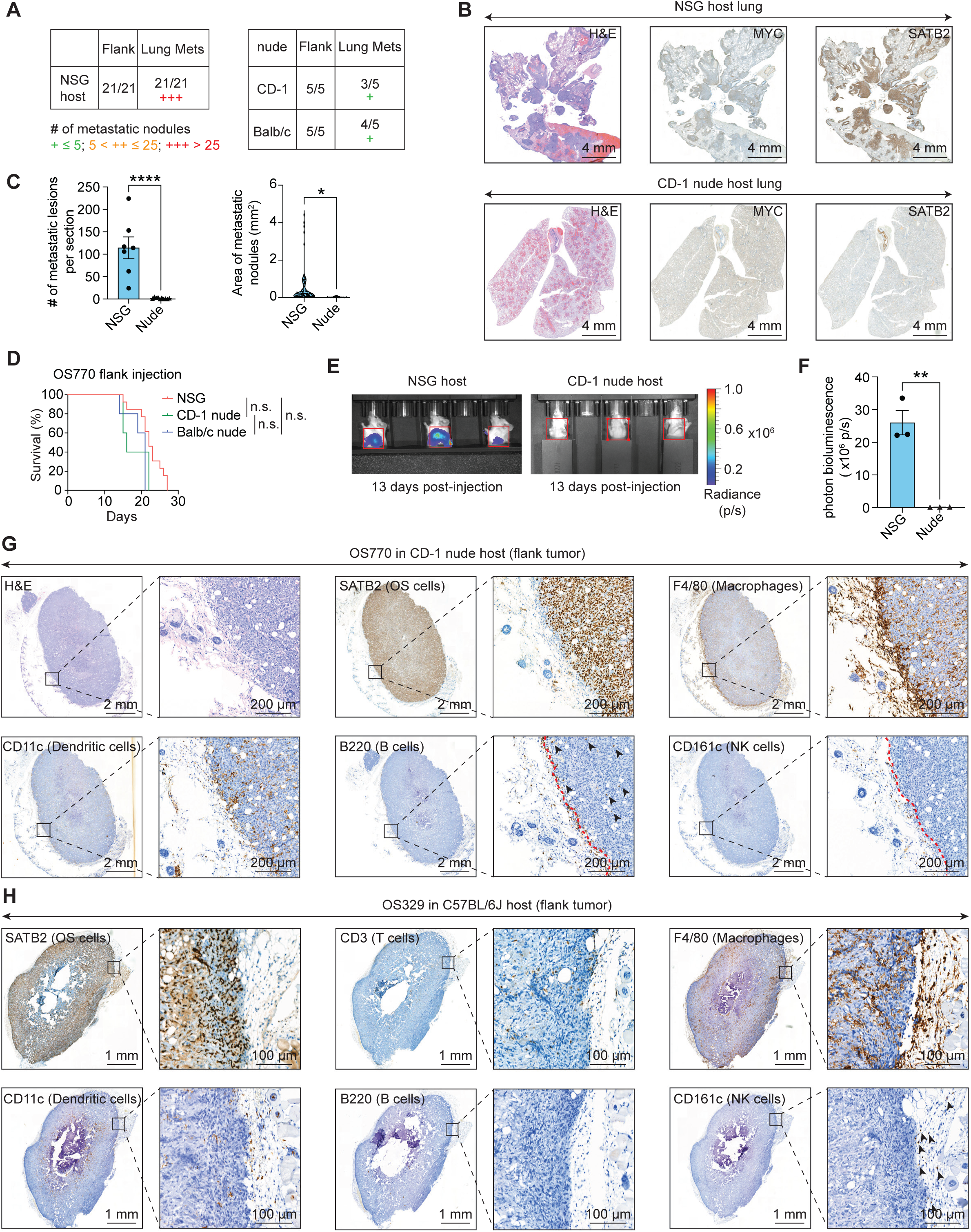
Subcutaneous implantation of OS770 induces lung metastases in immunocompromised NSG mice. (A) Quantification criteria and summary of OS pulmonary metastasis following subcutaneous flank injection of OS770 cells into independent cohorts of recipient mice. (B) NSG mice developed widespread MYC/SATB2-positive metastatic lung tumors, whereas nude mice developed limited micro-metastases that were detected in histological sections at the humane endpoint (noted in A). (C) Quantification of the number of lung metastatic lesions in NSG (n=7) and nude host (n=10), and the mean area of metastatic nodules (mm^2^/nodule). Data were shown as means ± SDs; statistical analysis was performed using the Student’s unpaired t-test: \**p* < 0.05, \*\*\*\**p* < 0.0001. (D) Survival curve for NSG (red, n=13) mice, CD-1 nude (green, n=5), and Balb/c nude (blue, n=5) mice with subcutaneous injection of OS770 into the flank. The humane endpoint was defined by the size of the flank tumor (<2,000 mm^3^) and respiratory difficulty. Statistical analysis was performed using the log-rank test: n.s. *p* > 0.05. (E) Bioluminescence detection was used to trace the luciferase-labeled OS770 cells in NSG and CD-1 nude mice at 13 days post-tumor implantation. Signals associated with lung metastases were captured while a shield blocked flank tumor signals. (F) Quantification of bioluminescence signals in the lungs of recipient mice. Data were shown as means ± SDs; statistical analysis was performed using the Student’s unpaired t-test: \*\**p* < 0.01. (G) Flank OS tumors were collected from CD-1 nude mice 13 days post-OS770 implantation. Serial sections were stained with H&E and different immune cell markers, including F4/80 (macrophages), CD11c (dendritic cells), B220 (B cells), and CD161c (NK cells). Boxes indicate the distribution of immune cells at the tumor-tissue junction and those infiltrated into the tumors. (H) Flank OS tumors were collected from C57BL/6J mice 41 days post-OS329 implantation. Serial sections were stained with SATB2 and different immune cell markers, including CD3 (T cells), F4/80 (macrophages), CD11c (dendritic cells), B220 (B cells), and CD161c (NK cells). Boxes indicate the distribution of immune cells at the tumor-tissue junction and those infiltrated into the tumors.

We next sought to determine which immune cell population might restrict OS metastasis in T cell-deficient nude mice. Immunohistochemistry revealed that macrophages (F4/80^+^) and dendritic cells (CD11c^+^) extensively infiltrated OS770 flank tumors in CD-1 nude mice, whereas B cells (B220^+^) and NK cells (CD161c^+^) were mainly marginalized to the tumor-host tissue boundary; no obvious CD161c^+^ NK staining was detected within the flank tumor itself (**Fig. 2G**). In the OS329-injected syngeneic model, some T cell infiltration (CD3^+^) was observed in C57BL/6J host flank tumors (**Fig. 2H**) in animals that survived somewhat longer than the NSG cohort (**Supplementary Fig. S4B**). B cells and NK cells were uniformly excluded at the periphery of tumor-host tissue junctions (**Fig. 2H**). In short, innate immune cells appeared to play a critical role in controlling metastasis, despite minimal effects on primary tumor growth.

### NK cell depletion promotes lung metastasis

The stage(s) at which innate immune cells prevent pulmonary metastasis could depend on restricting the migration of cells from the primary tumor, limiting hematogenous dissemination, and/or playing a critical defensive role for immune cells within the lung itself. To bypass the initial steps of metastasis, we applied an accelerated disease model by injecting luciferase-marked OS770 cells into the tail veins of recipient mice and tracking tumor migration by imaging (**Fig. 3A**). Intravenous tail vein injection of 3 × 10^6^ luciferase-labeled OS770 cells resulted in their prompt pulmonary trapping in both NSG and CD-1 nude hosts as early as one hour post-injection. Strong signals were still observed in the lungs of NSG mice 24 hours later, whereas no detectable signal was observed in CD-1 nude mice (**Fig. 3B**). Immunohistochemical staining for OS marker SATB2 revealed that numerous single OS cells were observed throughout the lungs of NSG mice at one hour after tail vein injection. Within 24 hours, these single cells had expanded into small metastatic colonies. In contrast, these disseminated OS cells were efficiently eliminated from the lungs of CD-1 nude mice, preventing colony formation (**Fig. 3C**). Subsequently, NSG mice exhibited massive pulmonary tumor burdens, necessitating their sacrifice 11-12 days after tumor engraftment. Strikingly, no metastases were found in the lungs of CD-1 nude hosts at the same time (**Fig. 3D and 3E**). Collectively, these data suggest that pulmonary innate immune cells eliminate tumor cell persistence in the lung.

**Fig. 3.**
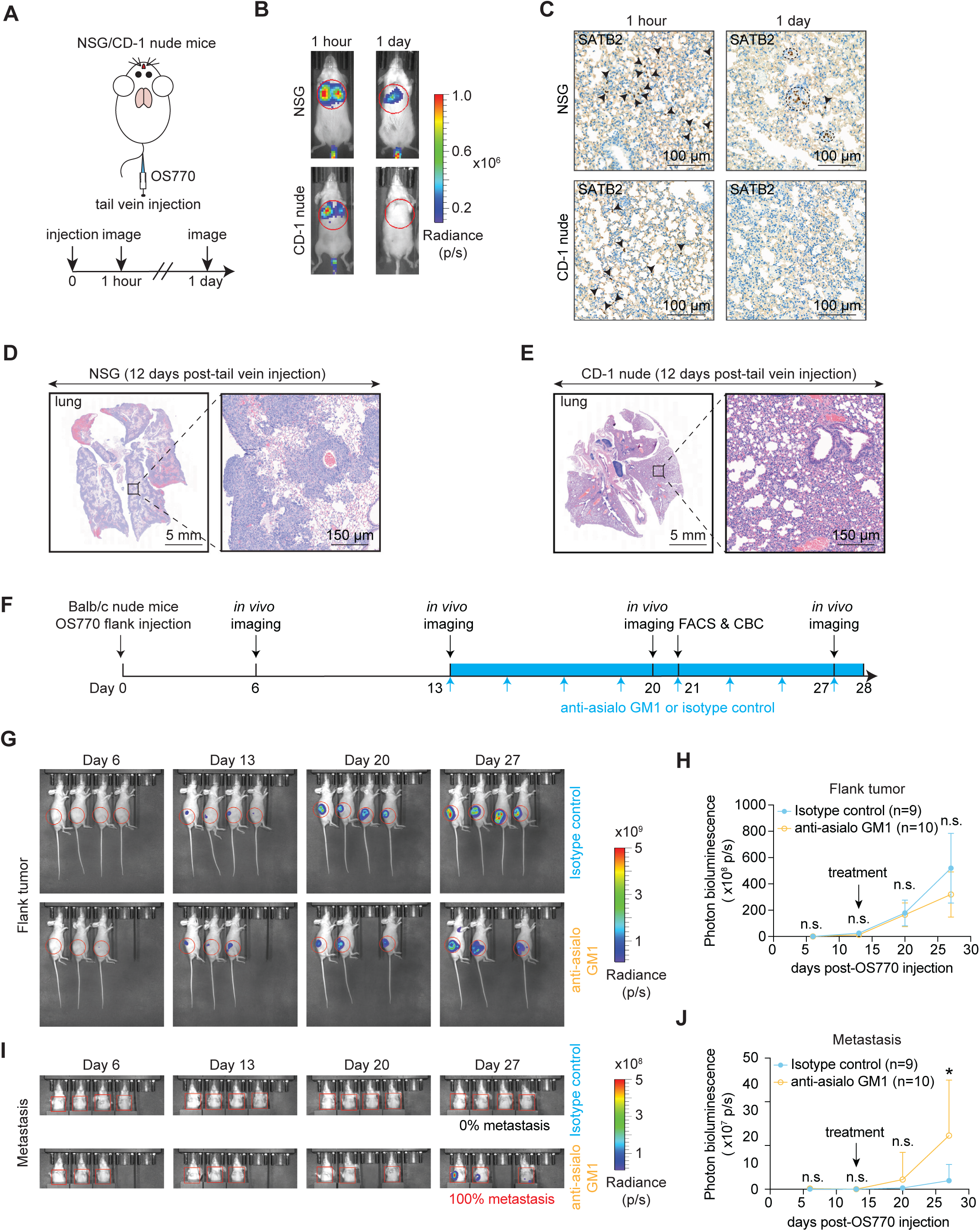
NK cells prevent OS lung metastasis in T cell-deficient nude mice. (A) Schematic of intravenous injection of 3× 10^6^ luciferase-labeled OS770 cells in NSG and CD-1 nude mice. The migration trajectory of OS cells was monitored by bioluminescent imaging at 1 hour and 1 day post-injection. (B) Representative whole-body bioluminescence images of NSG mice and CD-1 nude mice at one hour and one day after tail vein injection of 3 × 10^6^ OS770 cells. (C) Immunohistochemistry staining of SATB2 in the lungs of NSG and CD-1 nude mice at 1 hour or 1 day post intravenous injection of OS770 cells. Single tumor cells are labeled with arrowheads, and micro-metastatic colonies are labeled with circles. (D) Representative histopathological H&E-stained sections of the lung at 12 days after tail vein injection of 3 × 10^6^ OS770 cells in NSG mice. (E) Representative histopathological H&E-stained sections of the lung at 12 days after tail vein injection of 3 × 10^6^ OS770 cells in CD-1 nude mice. (F) Schematic diagram of the NK depletion protocol in Balb/c nude mice bearing flank OS tumors. Mice were subcutaneously injected with 3 × 10^6^ OS770 cells into the flank. About 13 days post-injection, mice were treated with either 50 μg of anti-asialo GM1 or an isotype control every other day until the humane endpoint. Tumor growth and lung metastasis were monitored by bioluminescence imaging on days 6, 13, 20, and 27. NK cell depletion efficiency in peripheral blood or the spleen was assessed by flow cytometry at one week after NK cell depletion and at the humane endpoint. The blood cell population was monitored by complete blood count (CBC). (G) Bioluminescence images of Balb/c nude mice bearing flank OS tumors and administered with isotype control antibody (top panels) or anti-asialo GM1 (bottom panels). (H) Quantification of signal intensity in flank OS tumors. Data from two independent animal cohorts were combined for statistical analysis: n (isotype control) = 9 and n (anti-asialo GM1) = 10. Data were shown as means ± SDs; statistical analysis was performed using the Student’s unpaired t-test at each parallel time point: n.s. *p* > 0.05. (I) Bioluminescence images of lung metastasis in Balb/c nude mice as shown in (G). (J) Quantification of signal intensity of lung metastasis. Data from two independent animal cohorts were combined for statistical analysis: n (isotype control) = 9 and n (anti-asialo GM1) = 10. Data were shown as means ± SDs; statistical analysis was performed using the Student’s unpaired t-test at each parallel time point: n.s. *p* > 0.05, \**p* < 0.05.

We elected to focus on NK cells, based on studies elsewhere that the secretome of drug-induced, senescent, primary pulmonary adenocarcinomas recruited cytotoxic NK cells rather than T cells to control tumors (27). Preliminary testing demonstrated that intraperitoneal injection of 250 μg anti-mouse NK1.1 neutralization antibody twice a week in C57BL/6J and CD-1 nude mice, or a single intraperitoneal injection of 50 μg anti-asialo GM1 administered two days prior in Balb/c nude mice, could efficiently deplete NK cells, as measured by rapid decreases of the NKp46^+^ population in spleens as determined by flow cytometry analysis (**Supplementary Fig. S6A-S6F**).

We first interrogated the role of NK cells in limiting OS lung metastasis in the context of an intact immune system. Immunocompetent C57BL/6J mice were treated with an anti-mouse NK1.1 antibody or an isotype control antibody twice weekly before subcutaneous injection of 3 × 10^6^ OS329 cells into the flank, with continuous administration of anti-mouse NK1.1 or isotype control antibody until the humane endpoint (**Supplementary Fig. S7A**). NK depletion in C57BL/6J mice showed limited effects on flank tumor development (**Supplementary Fig. S7B-S7C**). Complete blood counts and flow cytometry demonstrated that long-term administration of anti-mouse NK1.1 antibodies had minimal effects on circulating cell populations but specifically depleted NK cells. (**Supplementary Fig. S7D-S7G**). At the humane endpoint due to extensive flank tumor burdens, no visible lung metastases were observed during dissection, but histopathological sections identified micro-metastases that emerged in the lungs and axial lymph nodes of mice treated with anti-mouse NK1.1, but not in mice that received isotype control antibodies (**Supplementary Fig. S7H-S7J**).

To extend these findings in a T cell-deficient immune setting, Balb/c nude mice were subcutaneously transplanted with luciferase-labeled OS770 cells into the flank. When flank tumors reached approximately 300 mm^3^ in size 13 days after tumor engraftment, mice were treated with an anti-asialo GM1 neutralizing antibody or an isotype control every other day until the humane endpoint (**Fig. 3F**). Weekly bioluminescence imaging revealed that NK depletion had modest effects on flank tumor growth **(Fig. 3G and 3H)** but promoted lung metastasis in Balb/c nude mice (**Fig. 3I and 3J**). Flow cytometry and complete blood counts confirmed that anti-asialo GM1 treatment induced consistent NK depletion in both peripheral blood and spleen, with limited effects on other blood cell populations (**Supplementary Fig. S8A-S8D**). Microscopic review of histopathologic sections revealed extensive tumor burdens in the lungs of nude mice following NK depletion, as well as other organs, including the liver, lymph node, and adrenal gland, whereas only a few micro-metastatic lesions were found in mice treated with an isotype control antibody (**Supplementary Fig. S8E-S8H**).

To independently assess the protective effect of NK function in controlling lung metastasis, a second treatment regimen was employed. NK cells were pre-depleted for two days in Balb/c nude mice, followed by transplantation of luciferase-labeled OS770 cells into the flank (Day 0), with continuous injection of the anti-asialo GM1 neutralizing antibody or an isotype control antibody at two-day intervals until the humane endpoint was reached (**Supplementary Fig. S9A**). Again, metastatic signals emerged in all Balb/c nude mice treated with an anti-asialo GM1 neutralizing antibody, with minimal effects on flank tumor growth (**Supplementary Fig. S9B-S9E**). The NK populations in spleen and peripheral blood were consistently depleted following anti-asialo GM1 administration throughout the entire treatment regimen without affecting other blood cell populations (**Supplementary Fig. S9F-S9I**). Histopathological review following NK depletion in Balb/c nude mice confirmed the presence of extensive metastatic nodules in the lungs (**Supplementary Fig. S9J and S9K**), livers, lymph nodes, kidneys, and adrenal glands, with variable penetration from 20 to 100% in each distal organ (**Supplementary Fig. S10A and S10B**), phenocopying metastatic spread in NSG mice.

Given that NK cells control metastasis in our OS mouse model, we sought to investigate the clinical relevance of this effect. To this end, we surveyed publicly available data from human OS clinical trials in which patient tumors were evaluated for gene expression before receiving therapy (see **Supplementary Methods**). Indeed, higher expression of the NK cell surface marker *NCR1* significantly correlates with better survival outcomes in OS patients conventionally treated with surgery and chemotherapy (**Fig. 4A-4C**). To further evaluate NK function in OS patients, activity score was generated by combining 22 NK cell signature gene expression, including NK cell surface identity markers (*NCAM1/CD56* and *FCGR3A/CD16*), activating/inhibitory receptors (*NCR1, NCR3, KLRK1, KLRC1/2,* and *KLRD1*), cytotoxic machinery (*PRF1, GZMB/K/H, GNLY,* and *NKG7*), effector cytokines (*IFNG* and *XCL1/2*), signaling adaptors (*TYROBP, FCER1G,* and *CD247*), and chemokine receptors (*CXCR3* and *CX3CR1*). The results revealed that higher NK signature scores were notably associated with improved survival across three independent cohorts (**Fig. 4D**), highlighting a pivotal role for NK cells in OS.

**Fig. 4.**
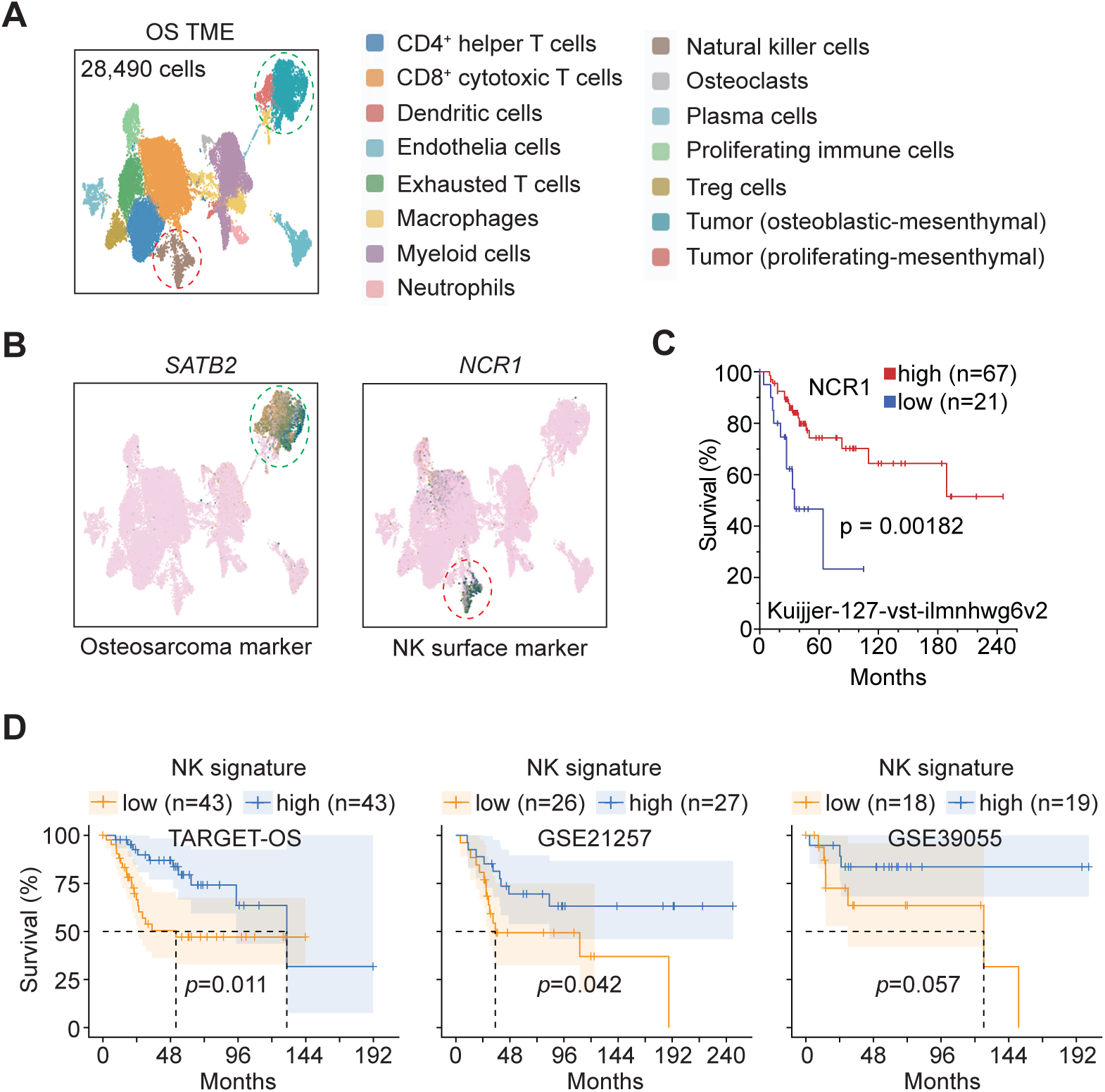
High expression of NK cell signature genes correlates with improved survival in OS cohorts. (A) Single-cell RNA-seq data were obtained from the publicly available OS dataset of Sid Sijbrandij (osteosarc.com), which comprises longitudinal tumor biopsies from a single patient with recurrent spinal osteosarcoma. A total of 28,490 OS tumor cells and other cell types within the tumor microenvironment (TME) were processed and stratified into 15 distinct populations based on signature gene expression. (B) OS tumor cells (labeled as green circle) highly express *SATB2*, and NK cells (labeled as red circle) highly express the NK surface marker gene *NCR1*. (C) High expression of the NK surface marker gene *NCR1* correlated with better overall survival outcomes in the Kuijjer-127-vst-ilmnhwg6v2 dataset. Patients were stratified into high (red, n = 67) and low (blue, n = 21) *NCR1* expression groups using an automated scan cut-off for the log-rank test. (D) NK cell signature gene expression [a cluster of surface identity markers (*NCAM1/CD56, FCGR3A/CD16*), activating/inhibitory receptors (*NCR1, NCR3, KLRK1, KLRC1/2, KLRD1*), cytotoxic machinery (*PRF1, GZMB/K/H, GNLY, NKG7*), effector cytokines (*IFNG, XCL1/2*), signaling adaptors (*TYROBP, FCER1G, CD247*), and chemokine receptors (*CXCR3, CX3CR1*)] and survival outcomes across the TARGET-OS cohort, GSE21257 cohort, and GSE39055 cohort. Patients were stratified into high (blue) and low (orange) NK signature gene expression groups using a median cut-off for the log-rank test.

### An unbiased screen for genes that regulate the OS secretome and pulmonary metastasis

To explore the molecular mechanisms underlying metastasis in OS cells, we employed an unbiased loss-of-function CRISPR screen to identify genes that enable OS770 cells to evade NK cell surveillance and metastasize to the lungs in nude mice. Such a screen would likely identify genes whose loss might enhance OS metastasis through a variety of mechanisms, although, for our purposes, we hoped to focus on targeted genes encoding activating NK receptor ligands. Considering the near-tetraploid nature of OS cells, we optimized genome-editing tools by establishing OS cell lines stably expressing Cas9 (OS770^Cas9^)(**Supplementary Fig. S11A and S11B**), which, like parental OS770 cells, formed lethal metastatic OS tumors in recipient NSG mice (**Supplementary Fig. S11C and S11D**). As proof of principle, an optimized construct expressing a single-guide RNA (sgRNA) was directed to *Myc* successfully in OS770^Cas9^ cells, resulting in impaired cell viability due to significant knockdown of MYC protein expression (**Supplementary Fig. S11E-S11G**).

To screen for candidate “metastasis-associated genes” using CRISPR, a whole-genome Brie sgRNA library targeting 19,674 mouse genes (about 4 sgRNAs per gene)(28) was used to infect OS770^Cas9^ cells, followed by their subcutaneous transplantation into the CD-1 nude mice. As a negative control, OS770^Cas9^ cells carrying the same pooled sgRNA library were transplanted into NSG hosts in which lung metastasis would likely occur regardless of perturbation of genes affecting the OS secretome (**Fig. 5A**). To facilitate enrichment of metastatic cells from the lung, lung tissues were collected for tumor cell dissociation and subsequent culture (**Fig. 5A; Supplementary Fig. S12A**). Over a seven-day expansion period, normal lung cells stopped growing, whereas metastatic tumor cells selectively and rapidly expanded (**Supplementary Fig. S12B and S12C**). These sgRNA-targeted tumor cells likely had escaped immune surveillance in nude mice, presumably because key genes that confer metastasis resistance were disrupted.

**Fig. 5.**
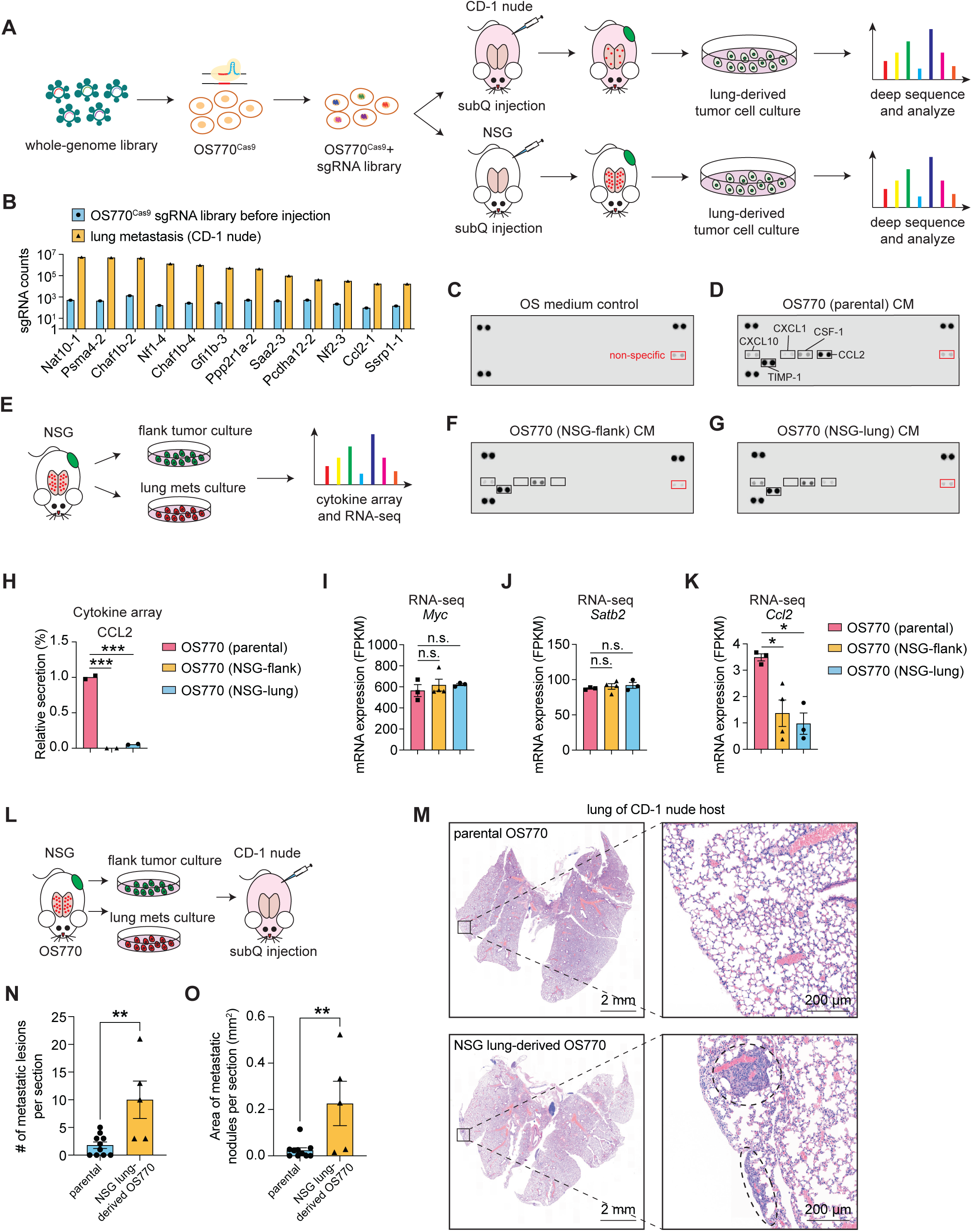
Genome-wide CRISPR screen identifies essential genes controlling lung metastasis. (A) Schematic diagram of a whole-genome loss-of-function CRISPR screen in OS770 cells to identify essential genes controlling lung metastasis. CD-1 nude and NSG mice were used as transplantation recipients in parallel to identify immune-relevant candidates. (B) MAGeCK analysis was conducted in sgRNA library-induced lung metastatic cells from CD-1 nude mice compared to parental library cells. The sgRNAs with a count enrichment >100-fold were listed. Each gene was targeted by four independent sgRNAs; the number following the gene name indicates the individual sgRNA in the library. (C) Cytokine array analysis was used to unbiasedly define and quantify secreted cytokines/chemokines from the OS tumor cells in culture. Basal OS medium for cytokine array was used as a negative control for (D), (F), and (G). (D) Cytokine array revealed that OS770 cells secrete CXCL10, CXCL1, M-CSF, CCL2, and TIMP-1 into the cultured medium (CM). (E) Schematic diagram of recovering NSG mice flank- and lung-derived OS tumor cells in tissue culture and subsequent profiling by cytokine array and RNA-seq. (F) Cytokine array profiling of the NSG flank tumor-derived OS770 cells. (G) Cytokine array profiling of the NSG lung tumor-derived OS770 cells. (H) Quantification of CCL2 secretion levels shown in (D), (F), and (G). (I) *Myc* mRNA expression levels in parental OS770 cells and OS770 cells derived from flank and lung tumors in NSG mice. (J) *Satb2* mRNA expression levels in parental OS770 cells and OS770 cells derived from flank and lung tumors in NSG mice (K) *Ccl2* mRNA expression levels in parental OS770 cells and OS770 cells derived from flank and lung tumors in NSG mice. (L) Schematic of recovering NSG mice flank and lung-derived OS tumor cells in tissue culture and subsequent subcutaneous implantation in CD-1 nude mice. (M) Images of H&E-stained lung sections from CD-1 nude mice subcutaneously injected with parental OS770 cells or NSG lung-derived OS770 cells. (N) Quantification of the number of pulmonary metastatic nodules per section. (O) Quantification of the total area of pulmonary metastatic nodules per section. In (H-K), (N), and (O), data were shown as means ± SDs; statistical analyses were performed using the Student’s unpaired t-test: n.s. *p* > 0.05, \**p* < 0.05, \*\**p* < 0.01, \*\*\**p* < 0.001. Gene expression values were collected by whole-transcriptome RNA-seq.

To identify targeted genes, genomic DNA from cultured metastatic tumor cells and from the input population before transplantation was extracted and amplified using primers flanking the sgRNA expression cassette, followed by deep sequencing and computational (MAGeCK) analysis (29). Only 12 sgRNAs showed >100-fold representation in lung metastatic cells from the CD-1 nude host compared with parental cells (**Fig. 5B**). Except for the chromatin remodeler *Chaf1b*, for which two individual sgRNAs were detected with >3,000-fold enrichment, one sgRNA against each of the other 10 genes (*Nat10*, *Psma4*, *Nf1*, *Nf2*, *Gfi1b*, *Ppp2r1a*, *Saa2*, *Pcdha12*, *Ssrp1*, and *Ccl2*) was overrepresented in lung tumors.

A very likely possibility is that CRISPR editing would target “metastasis genes” that play no direct role in the immune environment, exemplified by loss-of-function of tumor suppressor genes, such as *Nf1* and *Nf2*, both of which were enriched in our screen, and by transcription factors (e.g. *Gfi1b*)(30) and chromatin remodeling proteins (e.g. *Chaf1b*)(31) that contribute to tumorigenesis in other cancer models (**Fig. 5B**). To exclude such OS cell-autonomous metastasis mechanisms, a second round of screening was conducted by transplanting the sgRNA library-containing OS770 cells into NSG mice and measuring sgRNA enrichment in lung metastases. Again, *Nf1* and *Nf2* passed the statistical cut-off (FDR < 0.05 and ∣log_2_FC∣ > 2) and were significantly enriched in lung tumors compared to flank tumors (**Supplementary Fig. S13A**). When comparing flank tumors with parental library-containing cells before injection, *Nf1* and *Nf2* were the only two hits (**Supplementary Fig. S13B and S13C**). As expected, disruption of *Nf1* alone in OS770 cells significantly augmented pulmonary tumor burdens in CD-1 nude mice (**Supplementary Fig. S13D-S13I**). These data suggested that *Nf1 and Nf2* function as documented tumor suppressors in an NK-independent manner to promote cell-autonomous metastasis (32, 33).

Despite the variability and multiplicity of genetic copy-number variations in human OS, a majority of 108 patient OS samples and PDXs exhibit strikingly reduced expression of four COPII vesicle components, leading to secretory defects (34). These almost uniform findings focused our attention on the OS secretome, raising the possibility that a reduction in the production of particular chemokines and cytokines by tumor cells might inhibit the recruitment of innate immune cells that counter metastasis. Notably, disruption of *Ccl2* was identified in our unbiased screen as a contributor to metastasis in CD-1 nude mice, but not in metastasis-permissive NSG mice, where its putative role in enforcing NK-dependent immunity would be irrelevant (**Fig. 5B**). In turn, OS770 cell-conditioned medium contained readily detectable levels of CCL2. Two other chemokines, CXCL1 and CXCL10, and the macrophage colony-stimulating factor, CSF-1/M-CSF, were detected at lower levels (**Fig. 5C and 5D**). CCL2, CXCL1, and CXCL10 stimulate NK recruitment and function (35–37), whereas CSF-1 supports the viability and proliferation of macrophages (38). OS770 cells also secreted TIMP1, an inhibitor of matrix metalloproteases that, like NF1/2, might facilitate metastasis through NK-independent mechanisms (39). However, CRISPR targeting *Cxcl1* had a limited effect on the pulmonary metastasis in CD-1 nude mice, highlighting the specific role of CCL2 (**Supplementary Fig. S13J-S13L**). Notably, OS770 and OS329 cells recovered from the flank or pulmonary tumors in NSG mice significantly reduced CCL2 secretion (**Fig. 5E-5H; Supplementary Fig. S14A**), along with a slight decrease in CXCL10 and TIMP1, as well as a marginal increase in CSF-1 (**Supplementary Fig. S14B-S14E**). RNA-seq confirmed that *Ccl2* transcription decreased in NSG host-derived OS770 cells, with stable expression of both oncogenic *Myc* and the lineage-specific gene *Satb2* (**Fig. 5I-5K**), whereas the expression of *Cxcl1*, *Cxcl10*, *Timp1*, and *Csf1* remained unchanged or slightly upregulated (**Supplementary Fig. S14F-S14I**). Transplanting NSG lung-derived OS770 cells that exhibited low CCL2 secretion into metastasis-resistant CD-1 nude mice markedly increased pulmonary metastatic burden (**Fig. 5L-5O**), with no changes in flank tumor development (data not shown). Together, these findings imply a unique role of CCL2-CCR2 signaling in controlling OS pulmonary metastasis.

### Disruption of CCL2 secretion alone in OS cells facilitates pulmonary metastasis

Publicly available single-cell RNA-seq data from a cohort of 11 OS patients (GSE152048) revealed that *CCL2* is expressed by OS cells and broadly by other TME populations, whereas its receptor gene, *CCR2*, is mainly restricted to a subset of macrophages and NK cells (**Supplementary Fig. S15A-S15C**). Notably, in a similar cohort association analysis using patient-derived primary tumor samples, high *CCL2* expression correlates with improved clinical outcome in OS patients, but is not detected in other ovarian, breast, or liver cancers (**Fig. 6A and 6B**; see **Supplementary Methods**). To functionally test the role of tumor-derived CCL2 in preventing pulmonary metastasis, we used the CRISPR/Cas9 system to disrupt the *Ccl2* gene in OS770 cells, reducing CCL2 expression and secretion (**Fig. 6C and 6D**). CCL2 deficiency had limited effects on the transcriptome network (**Fig. 6E**). Flank tumor growth of sgCcl2 was comparable to that obtained with non-targeted sgRNA (sgNT) controls (**Fig. 6F and 6G**). Strikingly, in two independent cohorts, the *Ccl2* KO cells significantly promoted lung metastasis, yielding greatly increased numbers and areas of pulmonary metastatic nodules (**Fig. 6H-6K; Supplementary Fig. S15D and S15E**). The frequency of metastases in nude mice initiated by *Ccl2* disruption alone in OS cells was lower than that achieved in NSG or NK-depleted nude mice, implicating roles for other chemokines and cytokines in regulating innate immune tumor control. Nonetheless, our findings emphasize the role of NK cells specifically within the pulmonary microenvironment in limiting OS metastasis, the major cause of treatment failure in patients, and point to a specific NK signaling node whose enhancement might augment effective therapy.

**Fig. 6.**
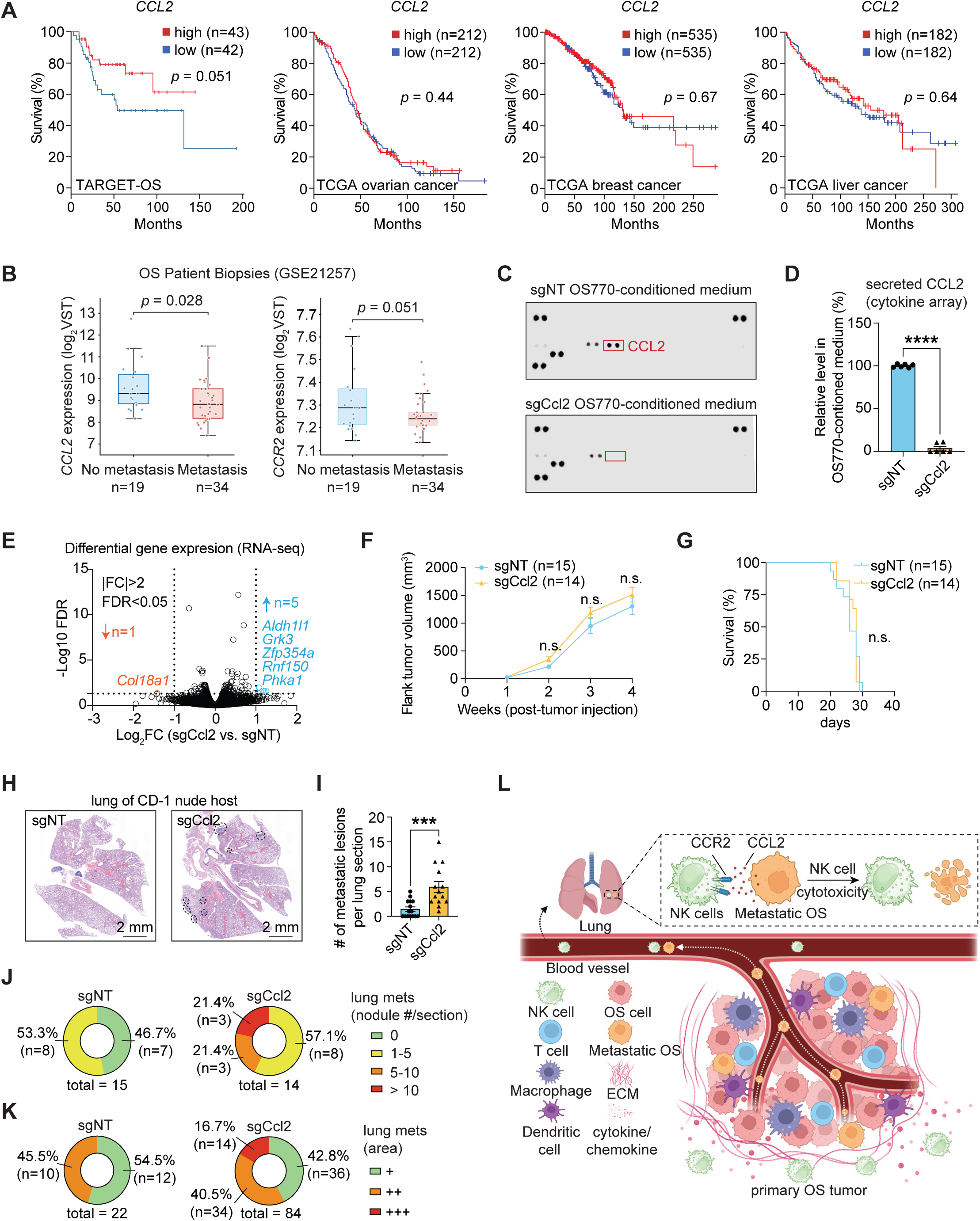
OS-derived CCL2 is pivotal to pulmonary metastasis. (A) Kaplan-Meier survival analysis of *CCL2* expression and its association with survival outcomes across different tumors, including the TARGET-OS cohort, the TCGA ovarian cancer cohort, the TCGA breast cancer cohort, and the TCGA liver cancer cohort. Patients were stratified into high (red) and low (blue) *CCL2* expression groups using a median cut-off for the log-rank test. (B) Transcriptomic gene expression data of 53 OS patients’ biopsies were downloaded from GSE21257. Patients were stratified into non-metastasis and metastasis groups based on clinical status and compared for *CCL2* and *CCR2* expression levels. Data were shown as means ± SDs; statistical analyses were performed using a Welch two-sample t-test. (C) Cytokine array confirmed significantly reduced CCL2 secretion in *Ccl2* KO OS770 cells. (D) Quantification of secreted CCL2 in sgNT and sgCcl2 OS770 cells. Data were shown as means ± SDs; statistical analysis was performed using a Student’s unpaired t-test. (E) Volcano plot of RNA-seq data identified differential gene expression in sgCcl2 versus non-targeted sgRNA (sgNT) OS770 cells. (F) The tumor growth curves in CD-1 nude mice bearing flank tumors induced by sgNT or sgCcl2 OS770 cells. Data were shown as means ± SDs; statistical analysis was performed using a Student’s unpaired t-test at each parallel time point, n.s. *p* > 0.05. (G) The survival curves of CD-1 nude mice bearing flank tumors induced by sgNT or sgCcl2 OS770 cells. Statistical analysis was performed using the log-rank test, n.s. *p* > 0.05. (H) Representative histopathological H&E-stained lung sections from the CD-1 nude mice bearing subcutaneous OS770 flank tumors expressing either sgNT or sgCcl2. (I) Statistics of the number of lung metastatic nodules per section. Data were shown as means ± SDs; statistical analysis was performed using a Student’s unpaired t-test, \*\*\**p* < 0.001. (J) Pie charts showing the percentage of lung metastatic nodule numbers in each CD-1 nude mouse per section. No metastatic nodules (green); 1-5 metastatic nodules (yellow); 5-10 metastatic nodules (orange); >10 metastatic nodules (red). (K) Pie charts showing the percentage of lung metastatic nodule areas in each CD-1 nude mouse per section. + (green) area ≤ 0.01 mm^2^; ++ (orange) 0.01 < area ≤ 0.05 mm^2^; +++ (red) area > 0.05 mm^2^. In (F-K), data from two independent animal cohorts were combined for statistical analysis: n (sgNT) = 15, n (sgCcl2) = 14. (A) Schematic of CCR2-CCL2 signaling-mediated NK-OS crosstalk in controlling pulmonary metastasis. The primary OS tumor lesion establishes an extracellular matrix (ECM) that limits NK cell infiltration, whereas T cells, macrophages, and dendritic cells can infiltrate the TME but fail to control primary OS tumor development. Metastatic OS tumor cells disseminate to distal organs, such as the lung, via the bloodstream. In the lung, metastatic OS cells secrete a suite of chemokines, including CCL2, to recruit CCR2-expressing NK cells, thereby facilitating NK-mediated cytotoxicity that eliminates metastatic OS cells and prevents pulmonary metastasis [Created in BioRender. Yang, J. (2026) https://BioRender.com/xsbvh0r].

## Discussion

The overarching goal of OS preclinical research is to discover effective therapies that might be translated to the clinic and benefit patients. To this end, reliable model systems can be indispensable tools for studying OS development, metastasis, and responses to therapy, and indeed, many investigators have stressed the limitations of, and the need for, new pre-clinical OS models (11, 40). Widely inconsistent results have been obtained across human and mouse OS cell lines, with only some metastasizing in immunodeficient mice, even after intravenous inoculation. Recently, orthotopic human PDX models have provided more reproducible avenues for OS metastasis research (8, 41, 42) but have not yet yielded actionable mechanistic insights or identified which host immune cells are most effective at combating disseminated disease. Reducing this complexity remains a challenge in devising effective therapeutic interventions. Leveraging our newly established mouse OS model, NK cell perturbation, and *in vivo* CRISPR screening, we revealed that CCL2 signaling-mediated NK-OS crosstalk plays a key role in restricting pulmonary metastasis (**Fig. 6L**).

The new mouse OS model described here combines enhanced Myc expression and *Trp53* loss, the most frequently mutated genes in OS, and mimics the cardinal features of OS human pathology, including widespread visceral metastasis with a distinct preference for the lung. When enriched *Trp53*-null, Myc-overexpressing GNPs were injected into the cerebral cortices of CD-1 nude mice, they routinely generated medulloblastomas and only rarely yielded other ill-defined tumors in a few animals (22, 23). However, when the Percoll-enriched *Trp53*-null GNP population was adapted to neurosphere culture and transduced with retroviral *Myc*, a minor population of mesenchymal tumor-initiating sphere cells selectively expanded and reproducibly induced OS when injected near the calvarium. Like their human counterparts, many *Trp53*-null OS cells lost tumor-suppressive *Pten* function; underwent whole-genome duplication to become near-tetraploid; and sustained many additional chromosomal abnormalities, including deletions, focal amplifications, and translocations. The emergence of primary OS tumors following injection of cultured donor sphere cells into the subcalvarial cortices of recipient mice arose after a 3-4 month latency, likely reflecting the selection of particular *Trp53*-null, Myc-amplified tumor-initiating mesenchymal progenitors that sustained further complex karyotype aberrations. Loss of p53 function alone contributes to the disruption of genome integrity, preventing the elimination or repair of damaged cells, blunting the apoptotic response to oncogene-induced stress, and transcriptionally engaging a broad network of tumor-suppressing target genes (43). However, widespread chromosomal instability in OS is a hallmark characteristic of chromothripsis, an ongoing process that occurs in at least 75% of human OS tumors but not in other cancers driven by *TP53* mutation (5, 6). Although induced by *TP53* inactivation and oncogene amplification, concurrent breakage-fusion-bridge cycles define a newly described loss-translocation-amplification (LTA) mechanism for OS not seen in many other *TP53*-null cancers (6). Our OS models indicated that both diploid and near-tetraploid karyotypes are stably maintained *in vivo* and in culture, suggesting that chromosome doubling is not strictly required for tumor initiation or maintenance of OS.

Comparative transplantation of OS cells into NSG versus nude mice revealed a requirement for innate immune surveillance in restraining pulmonary metastasis. Using intravenous tail-vein injection of OS cells to bypass the migration of tumor cells from primary tumors, OS cells rapidly accumulated in the lungs of both CD-1 nude and NSG mice, but were eradicated from CD-1 nude lungs while preferentially forming metastatic colonies in NSG recipients. Further antibody-based immune cell depletion experiments in Balb/c nude mice uncovered a principal role of NK cells, in particular, in limiting metastasis without significantly inhibiting the growth of primary tumors. Consistent with this, NK cell ablation in immunocompetent C57BL/6J or T cell-deficient Balb/c nude mice resulted in widespread micro-metastatic dissemination of OS cells from primary flank tumors to lymph nodes and other organs, including the liver, kidney, and adrenal gland. Notably, the more overt pulmonary metastasis following failure to eliminate colonizing tumor cells in the lungs points to a predominant protective role for NK cells within the lung microenvironment, as previously observed in lung adenocarcinomas (27), and recapitulates the frequent pulmonary metastasis in human OS.

In patients, administration of interleukin-2 (IL-2) augmented the absolute peak counts and activity of NK cells in the blood, which were positively correlated with improved OS clinical outcomes (44). Patient-derived chemotherapy-resistant OS cells were also vulnerable to elimination by IL-15-stimulated allogeneic and autologous NK cells (45). Higher expression of NK cell surface markers and NK network signature genes in patient biopsy samples also correlated with better clinical outcomes in patients conventionally treated for OS by surgery and chemotherapy. Ultimately, analysis of the OS secretome and CRISPR-based unbiased and targeted loss-of-function screens initiated with our OS cell lines revealed that CCL2 chemokine secretion by OS cells plays a key role in supporting NK defense against pulmonary metastasis, the most significant cause of treatment failure in OS patients. The major implication of this work is that enhancing NK activity in the lung, possibly with an NK CCR2 receptor agonist, could improve OS treatment. Targeting NK cells in the lung via tracheal vector delivery of ligands to activate pulmonary NK cells might be a focused approach to prevent OS metastasis in patients.

Despite our focus on metastasis control by the innate immune system, an unbiased CRISPR screen identified a suite of other potential “metastasis genes”. Among the 12 “hits” were tumor suppressors, *NF1* and *NF2*, and genes regulating transcription (*Gfi1b*), chromatin remodeling (*Chaf1b*, *Ssrp1*), proteasomal activity (*Psma4*), and RNA modification (*Nat10*), some of which have been implicated in promoting other cancers. Moreover, although we underscore the primacy of NK cells and CCL2 signaling in T cell-deficient models, we do not preclude the contribution of other cells within the TME to OS metastasis nor a role for adaptive immunity in fully immunocompetent animals. Indeed, whole-body depletion of CCL2 using a neutralizing antibody appeared to suppress OS metastasis in a previous study (46). In our system, OS tumor cell-specific disruption of CCL2 facilitates lung metastasis. These data suggest that the origin and dosage of CCL2 might be critical for controlling OS metastasis. Not surprisingly, antibody-mediated NK depletion had a more profound effect on metastasis than *Ccl2* disruption alone, even in T cell-deficient mice, indicating that other NK-OS signaling cascades likely play a role. OS cells secrete CSF-1 to maintain pro-tumorigenic M2-polarized macrophages, which represent the major cell type within the OS TME (10). In a broader context, cancer-associated fibroblasts (CAFs) and adaptive immune cells, including regulatory T cells, can further influence tumor growth and metastasis, yielding contrasting results across different models (10). Our institutional colleagues recently reported that redirecting B7-H3 CAR-T cells to chemokines expressed by OS cells enhanced their homing and anti-tumor activity in preclinical xenograft models, providing evidence supporting future CAR-T cell therapies (47, 48). Another clinical trial using NKG2D.zera-CART-NK cells and GD2-CART-T cells to treat human OS is currently open to recruit patients (NTC07211737). At the genome level, the complexity of nonuniform mutations coupled with chromothripsis and ongoing genomic instability in human OS poses a significant challenge for identifying additional drivers of disease progression and therapeutic gene targets. Nonetheless, reduced secretion of specific molecules from OS cells, due to downregulation of COPII proteins and independent of their mutational burden, underscores a general role of the OS secretome in engaging the innate immune system (34). Unveiling and targeting relevant signaling pathways, including the OS-CCL2-NK axis, offers a therapeutic approach to interfere with metastasis, the major cause of OS treatment failure.

## Methods

Previously used methods adapted with minor revisions for the present studies are described as **Supplementary Methods.**

### Animal husbandry and breeding

The immunocompromised mice, [CD-1 *nu/nu* (nude, stock 086; Charles River Laboratories), NSG mice (stock 005557; The Jackson Laboratory), Balb/c *nu/nu* mice (stock 194; Charles River Laboratories)], and immunocompetent mice C57BL/6J (stock 00664; The Jackson Laboratory) were used as tumor recipient mice at the age of 6-8 weeks. The C57BL/6J *Trp53*-null mice (stock 002101; The Jackson Laboratory) were used as a source of GNPs. For lineage tracing experiments, *Atoh1-Cre^ERT2^* (stock 7684; The Jackson Laboratory) and the dual fluorescent tdTomato-EGFP-Cre reporter allele (*mT/mG*; switch of tdTomato expression to EGFP after Cre recombinase; stock 007676; The Jackson Laboratory) were crossed into a *Trp53*-null background. *Atoh1-Cre^ERT2^*; *Trp53*-null males were crossed with *mT/mG*; *Trp53^+/-^* females to generate *Atoh1-Cre^ERT2^*; *mT/mG*; *Trp53*-null pups. At postnatal days 0 (P0) and 1 (P1), pups were injected intraperitoneally with 0.075 mg/g of Tamoxifen (stock: 5 mg/mL in corn oil) to induce Cre recombinase translocation to the nuclei. All animal experiments were approved by and conducted in accordance with St. Jude Children’s Research Hospital Animal Care and Use Committee guidelines, as required by the United States Animal Welfare Act and the National Institutes of Health’s policy to ensure proper care and use of laboratory animals for research.

### Vector construction

The MSCV retrovirus-based plasmid expressing mouse Myc-HA-miniAID-IRES-OsTIR1^F74G^-P2A-EGFP was generated using a two-step cloning strategy. First, mouse MSCV-Myc-IRES-luciferase was cut with BamHI and BstXI to remove the luciferase cDNA sequence. The OsTIR1^F74G^ (Ostir1 F and Ostir1 R primers) and P2A-GFP (P2A GFP F and P2A GFP R primers) fragments were amplified by PCR from pCDH-MND-OsTIR1^F74G^-P2A-EGFP-AID2-EF1a-RFP (#232800; Addgene) and XLone-Puro-Cas13d-EGFP-U6-BbsI (#155184; Addgene), respectively, and cloned to the digested MSCV-Myc-IRES backbone to make the intermediate MSCV-Myc-IRES-OsTIR1F74G-P2A-GFP plasmid. MSCV-Myc-IRES-OsTIR1^F74G^-P2A-EGFP was digested with EcoRI and BstBI to clone the HA and the miniAID tag (HA miniAID F and HA miniAID R) into the C-terminus of mouse Myc (Myc^HA-miniAID^) to generate the MSCV-Myc-HA-miniAID-IRES-OsTIR1^F74G^-P2A-EGFP plasmid. The NEBuilder HiFi DNA assembly kit was used for all cloning reactions according to the manufacturer’s recommendations (#E5520S; NEB). All PCR reactions used CloneAmp polymerase according to the manufacturer’s recommendations (#639298; Takara). Lentiviral pCDH-CMV-Myc-EF1-RFP was used to express wild-type (WT) Myc (Myc^WT^) in neural sphere cells. Lenti-Cas9-BSD (Blasticidin S) (#83480; Addgene) was used to express Cas9 in cells. To trace the sgRNA-targeted cells, the IRES-CFP cassette was inserted into the Lenti-Guide-Puro plasmid (#52963; Addgene) by In-Fusion cloning using CFP F and CFP R primers. The sgMyc (sgMyc F and sgMyc R) and sgCcl2 (sgCcl2-1 F, sgCcl2-1 R, sgCcl2-2 F, sgCcl2-2 R) oligonucleotides were synthesized (Thermo Fisher Scientific), annealed, and cloned into the Lenti-Guide-Puro-IRES-CFP construct between two BsmBI sites. All cloning primers are listed in **Supplementary Table S3**.

### Cell culture

Cerebella were isolated from 7-day-old mice of the desired genotype, dissociated with Trypsin treatment into a single cell suspension, and layered on a 35-60% Percoll gradient column as described (49). Cells, collected at the interface of 35-60%, were grown directly in Neurobasal medium (#21103049; Gibco) containing 1% penicillin/streptomycin (#15140163; Thermo Fisher Scientific), 2 mM L-glutamine (#G7513; Sigma), 1 × B27 (#175-04-044; Invitrogen), 1 × N2 (#175-02-048; Invitrogen), 30 ng/mL EGF (#AF-100-15; Peprotech), and 30 ng/mL FGF (#AF-100-18B; Peprotech) on an ultra-low-attachment surface. Cells from spheres were dissociated by Accutase (#A11105-01; Invitrogen) and infected with lentivirus co-expressing mouse Myc^WT^ and RFP or with retrovirus co-expressing mouse Myc^HA-miniAID^, OsTIR1^F74G^, and GFP. High Myc^WT^ or Myc^HA-miniAID^-expressing cells were sorted for the top 10% RFP^+^ or GFP^+^ population, respectively, and expanded in sphere culture. About 2 × 10^6^ cells, resuspended in 5 μL of Matrigel (356234; Corning), were orthotopically injected into the cortices of CD-1 nude mice as described (49). OS lines were obtained by dissociating the cranial bone tumors using the mouse tumor dissociation kit (#130-096-730; Miltenyi Biotec) following the manufacturer’s instructions. OS cells subcutaneously injected into NSG hosts developed flank tumors and lung metastases. Cells derived from flank tumors or lungs of host mice were isolated using the same protocol. Isolated cells were grown in SmGM-2 (#CC-3182; Lonza) for adherent cell expansion. The 293T cells were cultured in DMEM medium (#11965118; Gibco) containing 10% fetal bovine serum (FBS) (#SH30088.03; Hyclone), 2 mM L-glutamine, and 1% penicillin/streptomycin. Cells were routinely tested for mycoplasma contamination using a LookOut Mycoplasma PCR Detection Kit (#MP0035; Sigma) following the manufacturer’s instructions. All cells were cultured in a 37 °C incubator with a 5% CO_2_ atmosphere and 95% humidity.

### NK cell depletion

For Balb/c nude mice, anti-asialo GM1 (#014-09801; Fujifilm Irvine Scientific; 50 µg in 100 µL of 50% ddH_2_O and 50% RPMI) was injected intraperitoneally every other day. Rabbit IgG control (#011-000-003; Jackson Immuno Research Laboratories; 50 µg in 100 µL of 50% ddH_2_O and 50% RPMI) was used as a control. For C57BL/6J and CD-1 nude mice, anti-mouse NK1.1 (#NB0036; BioXCell; 250 µg in 100 µL PBS) was injected intraperitoneally twice weekly. IgG2a isotype control (#BP0085; BioXCell; 250 µg in 100 µL PBS) was used as a control. The NK cell depletion efficiency was assessed by flow cytometry. Unclotted peripheral blood was collected in EDTA anticoagulant. Splenocytes were collected in PBS containing 2% of FBS by mashing the spleen through a 70 µm strainer. Red blood cells were lysed in ACK hypotonic buffer (#3142792; Gibco). About 1 × 10^6^ splenocytes or blood cells were incubated in 200 µL of diluted anti-NKp46 conjugated with PE (1:200 in PBS containing 2% FBS, #137647, BioLegend) or conjugated with PerCP/Fire 806 (#137649; BioLegend; 1:200). The percentage of NKp46^+^ NK cells was determined by flow cytometry. Data analysis and presentation were performed using FlowJo software.

### CRISPR screening and data analysis

The whole-genome mouse CRISPR knockout pooled (Brie) library (28)(#73633; Addgene) was used. The Cas9-expressing OS770 cells were infected with the pooled sgRNA library at a low multiplicity of infection (M.O.I, ∼0.3) and selected with Blasticidin S and puromycin for 3 days. For the Brie sgRNA library screen, five CD-1 nude or NSG recipient mice were injected subcutaneously with 3 × 10^6^ OS770 cells carrying a pooled sgRNA library. Mice were euthanized at the humane endpoint, and the lungs were collected for metastatic tumor cells purification using the mouse tumor dissociation kit following the manufacturer’s instructions. Tumor cells were cultured under OS conditions with puromycin selection to expand to sufficient numbers. OS cells carrying the sgRNA library from one of five CD-1 nude mice and four of five NSG mice were successfully expanded. The genomic DNA was extracted using a PureLink Genomic DNA extraction Kit according to the manufacturer’s protocol. The sgRNA sequences were recovered by genomic PCR and submitted for deep sequencing on the MiSeq using single-end 150 bp reads (Illumina). The primer sequences used for cloning and sequencing are listed in the **Supplementary Table S3**. The raw FASTQ data were de-barcoded and mapped to the original sgRNA reference library. The differentially enriched sgRNAs were defined by comparing normalized counts between lung-derived tumor cells and parental bulk cells before injection. Normalized counts for each sgRNA were extracted and used to identify differentially enriched sgRNAs by DESeq2 (50), and the combined analysis was conducted by using the MAGeCK algorithm (29) as previously described.

## Data Availability

Raw sequencing data generated in this study have been deposited in the Gene Expression Omnibus (GEO). Specifically, the scRNA-seq data have been deposited under accession number GSE337056; the raw sequencing data for the CRISPR screen have been deposited under accession number GSE337055; and the raw sequencing data for the WGS have been deposited in the Sequence Read Archive (SRA) under PRJNA1483495. Raw data and experimental materials generated from this study are also available upon request to the corresponding author.

## Acknowledgment

We thank the members of the Li lab for their discussion and comments. We sincerely thank Sarah Robinson for mouse colony maintenance. We thank Drs. Xin Geng and Sathish Srinivasan for assistance with immunofluorescence staining. We deeply thank Dr. Liusheng He for assistance with flow cytometry analysis. We thank Teresa Santiago for the staining of the pathological sections. We gratefully acknowledge the staff of the Hartwell Sequencing Core Facility, the Flow Cytometry and Cell Sorting Shared Resource, the Vector Development and Production Laboratory, the Cytogenetics Laboratory, the Center for Applied Bioinformatics, the Animal Resource Center, the Comparative Pathology Core, and the Center for In Vivo Imaging and Therapy at St. Jude Children’s Research Hospital. This work was funded by an American Cancer Society Scholar Grant (RSG DMC-135487, to CL), V Foundation Scholar Grant (V2021-010, to CL), and American Lebanese Syrian Associated Charities (ALSAC, to CL). CJS, an HHMI emeritus, committed residual HHMI funds that support his continued research.

## Contribution Statement

Chunliang Li: Conceptualization, Data curation, Formal analysis, Funding acquisition, Project administration, Resources, Supervision, Visualization, Writing and editing the manuscript

Jifeng Yang: Data curation, Formal analysis, Investigation, Methodology, Project administration, Validation, Visualization, Writing and editing the manuscript

Frederique Zindy: Data curation, Formal analysis, Investigation, Methodology, Project administration, Validation, Visualization, Reviewing and editing the manuscript

Shaela Fields: Resources, Investigation, Methodology

Judith Hyle: Resources, Investigation, Methodology

Laura J. Janke: Methodology

Qianqian Li: Methodology

Beisi Xu: Resources, Formal analysis, Methodology

Qiong Zhang: Formal analysis, Methodology

Gang Wu: Resources, Formal analysis, Methodology

Ti-Cheng Chang: Formal analysis, Methodology

Monika Wierdl: Methodology

Lillian M. Guenther: Methodology

Linda M. Hendershot: Conceptualization

Martine F. Roussel: Resources, Supervision, Reviewing and editing the manuscript

Charles J. Sherr: Conceptualization, Funding acquisition, Supervision, Writing and editing the manuscript

