## Supplemental figures, legends, tables and methods. for "Chemokine-Dependent Natural Killer Cells Prevent Pulmonary Metastasis in a Newly Established Mouse Osteosarcoma Model"

**The combined supplementary file contains the following documents.**

Supplementary Figures S1-S15 and corresponding figure legends

Supplementary Tables S1-S3 and the corresponding table legends

Supplementary Methods

References

### Supplementary Figure S1

**A** SKY analysis (OS770, near-tetraploid)

70-75<4n>,XXYY,-3,-4,-7,-9,-10,dup(10),
-13,-13,der(14)t(14;19)x2,+del(?17)x2,
-19,-19[cp14]

73-74<4n>,XXYY,-3,-4,del(6),-7,-9,-10,dup(10),
-13,-13,der(14)t(14;19)x2,+del(?17)x2,
der(18)t(6;18),-19,-19[cp3]

75-78<4n>XXXY,+Y,-3,-4,-7,-9,dup(10),del(11),
+i(11)del(11),-13,-13,der(14)t(14;19)x2,
+del(?17)x2,-19,-19[cp3]

**B** SKY analysis (OS329, near-tetraploid)

69~74<4n>,XX,-9[11],-12[20],-12[20],
-13[20],-13[20],-14[20],-14[20],-18[13],
+1~4mar[20][cp20]

Annotation:
<4n> = indicates that all alterations are expressed in relation to the tetraploid level
del = deletion der = derivative dup = duplication i – isochromosome x2 = two copies
mar = 1-4 small marker chromosomes, but not all metaphases contain all markers

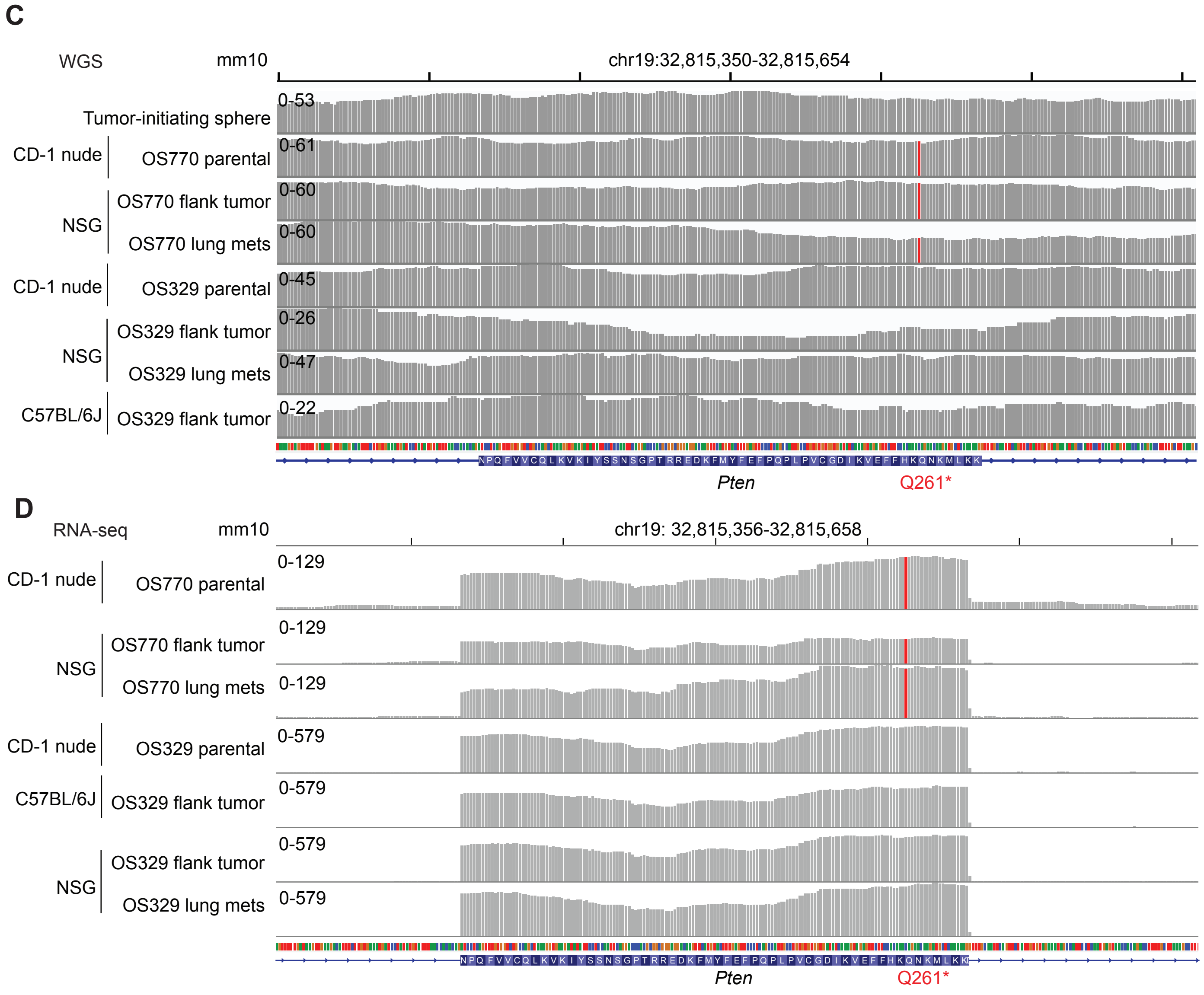

**Supplementary Figure S1. Established OS cells exhibit complex genomic rearrangements.**

(A) Fluorescence-labeled spectral karyotyping (SKY) analysis revealed chromosomal aberrations in OS770 cells.

(B) Fluorescence-labeled spectral karyotyping (SKY) analysis revealed chromosomal aberrations in OS329 cells.

(C) Whole-genome sequencing (WGS) revealed the nonsense mutation (Q261*) within the *Pten* locus in the OS770 cells.

(D) RNA sequencing (RNA-seq) revealed the nonsense mutation (Q261*) within the *Pten* locus in the OS770 cells.

St. Jude - Confidential

### Supplementary Figure S2

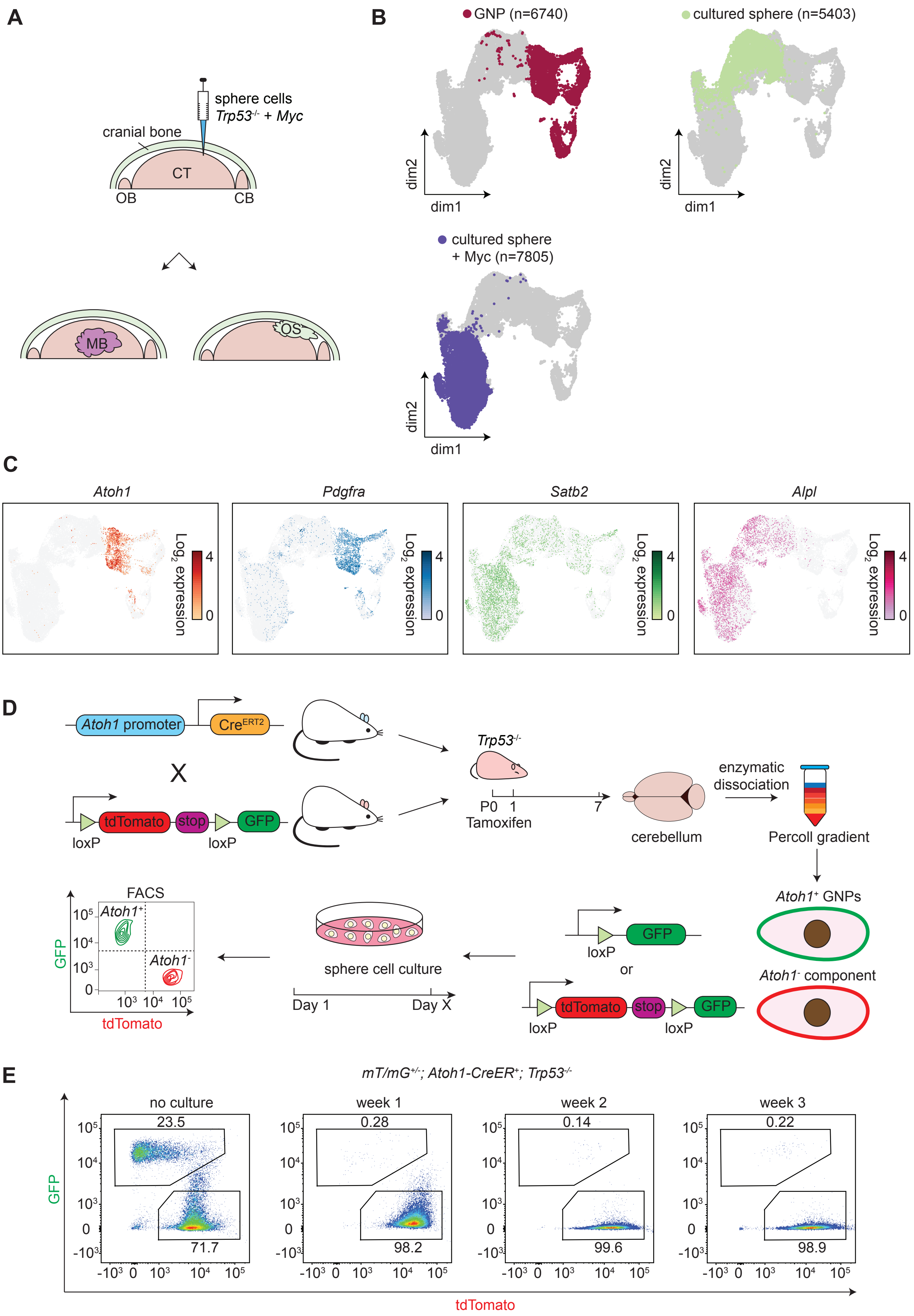

**Supplementary Figure S2. The Atoh1-positive granule neuron progenitor (GNP) population declines during in vitro sphere culture conditions.**

(A) Schematic diagram of two distinct tumor types (group 3 medulloblastoma, G3 MB and osteosarcoma, OS) originating from the *Trp53*-null sphere cells carrying ectopic Myc that were injected into the cortices of CD-1 nude mice.

(B) Freshly enriched GNPs (n=6,740, red), cultured sphere cells (n=5,403, green), and sphere cells with ectopic Myc (n=7,805, indigo) were collected for single-cell RNA-seq. Samples were sequenced individually and combined for Harmony integration analysis.

(C) The expression of *Atoh1* (GNP marker), *Pdgfra* (mesenchymal progenitor marker), *Satb2* and *Alpl* (osteoblast-associated marker) across different cell types.

(D) Schematic diagram of lineage tracing for Atoh1-positive GNPs. Male *Trp53*$^{-/-}$ mice carrying *Atoh1* promoter-driven tamoxifen-inducible recombinase Cre$^{ERT2}$ (*Atoh1-Cre*$^{ERT2}$) were crossed with *Trp53*$^{+/-}$ female mice harboring *mT/mG* reporter, generating *Trp53*$^{-/-}$ offspring that simultaneously carried one allele of *Atoh1-Cre*$^{ERT2}$ and the other allele of *mT/mG* reporter. Tamoxifen was administrated to neonatal pups on postnatal day 0 (P0) and P1 to induce Cre translocation to the nucleus. The pups were euthanized on P7, and the cerebella were isolated by enzymatic dissociation and Percoll gradient purification to enrich the GNPs fraction. The *Atoh1*-positive cells are genetically labeled with GFP due to Cre-mediated excision of the tdTomato-stop signal cassette, whereas *Atoh1*-negative cells retain tdTomato expression. This enriched GNPs fraction was cultured *in vitro* under sphere culture conditions. The proportions of GFP$^{+}$ and tdTomato$^{+}$ cell populations were monitored weekly by flow cytometry.

(E) Weekly flow cytometry of cultured sphere cells.

St. Jude - Confidential

### Supplementary Figure S3

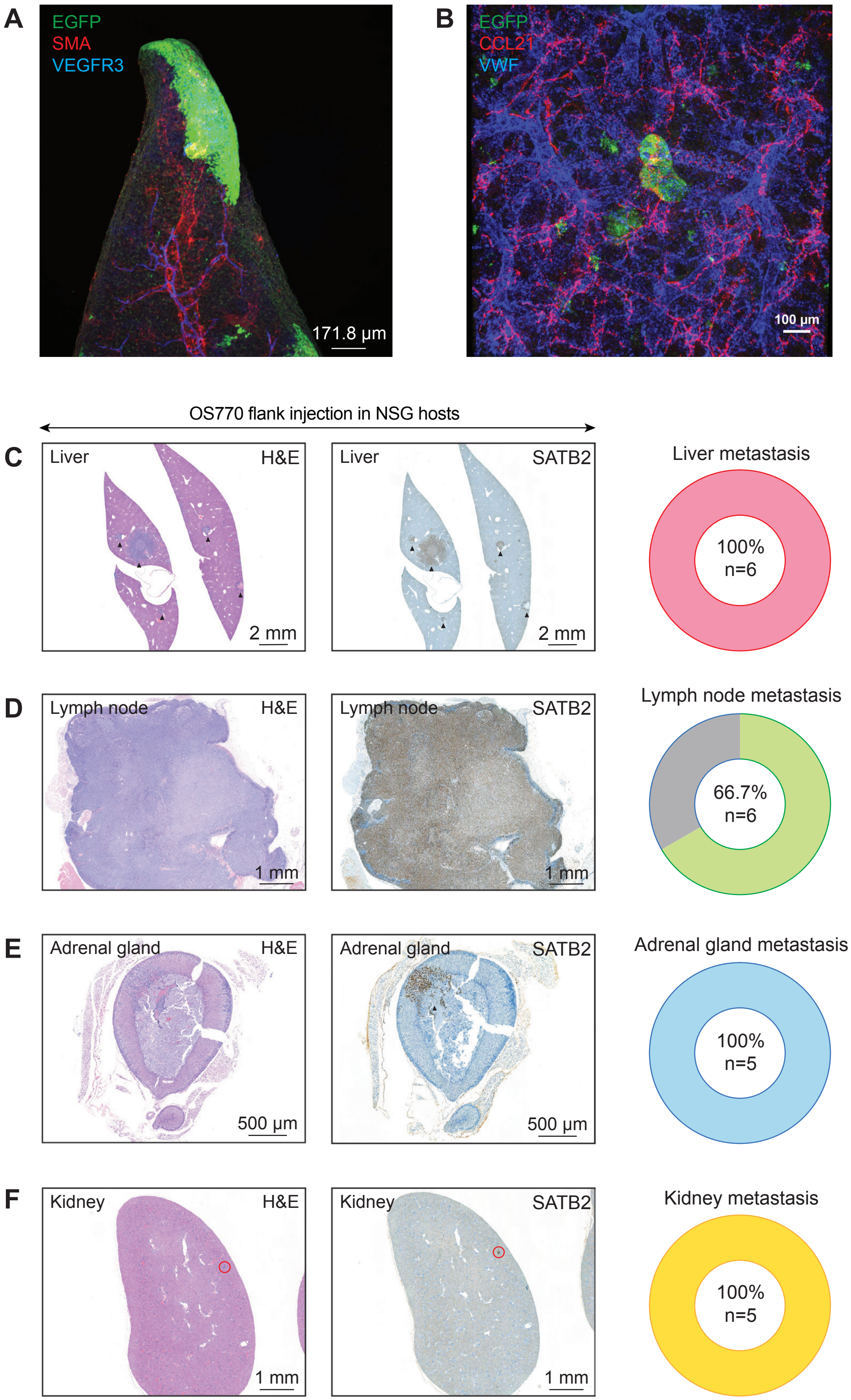

**Supplementary Figure S3. Subcutaneously injected OS770 metastasizes to distal tissues in NSG mice.**

(A) Representative immunofluorescence images showed that pulmonary metastatic OS cells cluster on the surface of the lung in NSG mice. The entire lung lobe was stained with anti-EGFP (OS cells; ectopic Myc expression cassette co-expressing GFP, labeled as green), anti-SMA (vascular smooth muscle cells, labeled as red), and anti-VEGFR3 (lymphatic endothelial cells, labeled as blue).

(B) Representative immunofluorescence images showed that pulmonary metastatic OS cells cluster in the parenchyma of the lung in NSG mice. The entire lung lobe was stained with anti-EGFP (OS cells, labeled as green), anti-CCL21 (lymphatic endothelial cells, labeled as red), and anti-VWF (vascular endothelial cells, labeled as blue).

(C-F) Representative histopathological sections and statistics for metastatic OS770 in the liver (C), lymph nodes (D), adrenal gland (E), and kidney (F).

St. Jude - Confidential

### Supplementary Figure S4

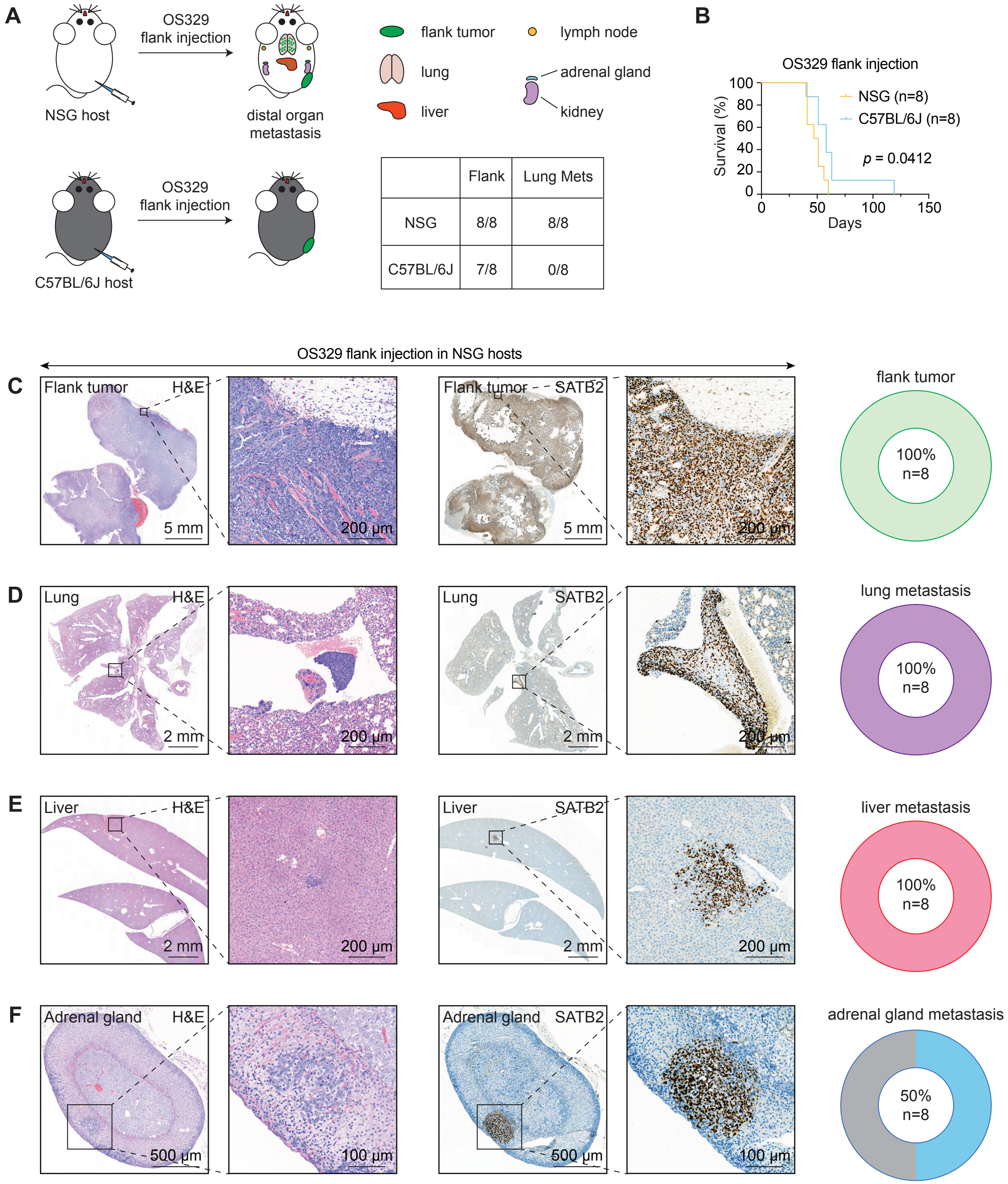


| | Flank | Lung Mets |
|---|---|---|
| NSG | 8/8 | 8/8 |
| C57BL/6J | 7/8 | 0/8 |

**Supplementary Figure S4. Subcutaneously injected OS329 metastasizes to distal organs in NSG hosts but not in immunocompetent C57BL/6J hosts.**

(A) $3 \times 10^6$ OS329 cells were subcutaneously injected into NSG hosts, which developed bony flank OS tumors and distal organ metastases. C57BL/6J immunocompetent mice developed flank tumors, but no metastases were detected.

(B) Survival curve of NSG and C57BL/6J mice bearing flank OS329 tumors. Statistical analysis was performed using the log-rank test.

(C-F) Representative histopathological sections and statistics for flank tumors (C) and metastases to the lung (D), liver (E), and adrenal gland (F) in NSG mice.

St. Jude - Confidential

### Supplementary Figure S5

**A**

OS770 NSG-flank

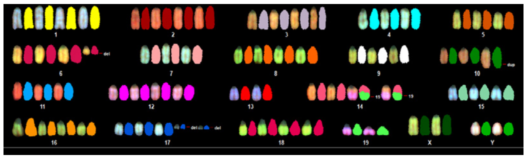

70~76, XXYY, der(Y)t(Y;6)(?D;?)[1], del(6)(?B)x1-2[17], del(10)(A1)[2], dup(10)(?)[17], -13[17], -13[17], der(14)t(14;19)(?C2;?B)x2[17], +der(?17)(?A2)x1-2[17],-19[17]-19[17] [cp17]

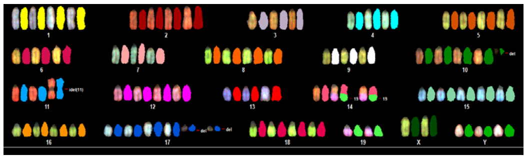

74~75, XXYY, del(10)(A1)[1], idel(11)[2], der(14)t(14;19)(?C2;?B)x2[2], +der(?17)(?A2)x1-2[2], -19[2], -19[2] [cp2]

**B**

OS770 NSG-lung

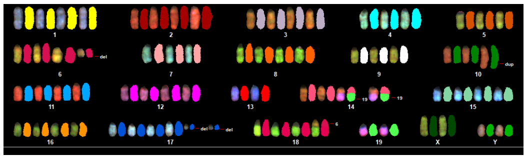

70~75, XXYY, del(6)(?B)[10], del(10)(A1)[2], dup(10)(?)[11], -13[11], -13[11], der(14)t(14;19)(?C2;?B)x2[11], +der(?17)(?A2)x2[11], der(18)t(6;18)(?;?)[10], -19[11]-19[10] [cp11]

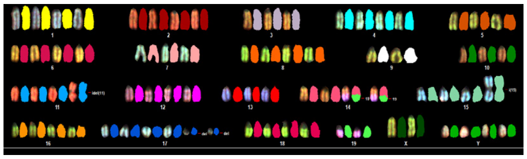

73~77, XXYY, i(1)[3], idel(11)[10], der(14)t(14;19)(?C2;?B)x2[10], i(15)[3], +der(?17)(?A2)x2[10], -19[10], -19[10] [cp10]

**C**

OS329 NSG-flank

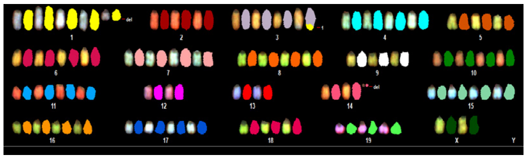

64~72, XX, -X/Y, +del(1)(?B), del(2)(A1), der(3)t(1;3)(?;?H), -9, +del(10)(A1)x1-2, -12, -12, -13, -13, -14, del(14)(A1), -18, -19[cp17]

**D**

OS329 NSG-lung

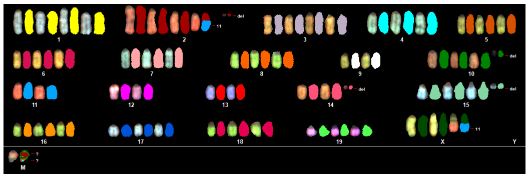

60~69, XX, der(?X)t(X;11)(?;?B1), +1, +der(2)t(2;11)(?B;?B1), +del(2)(A1), +3, del(9)(?A4), +del(10)(A1)x1-2, -11, -12, -13, del(14)(A1), +del(15)(?A2), der(?;?)[cp16]

**E**

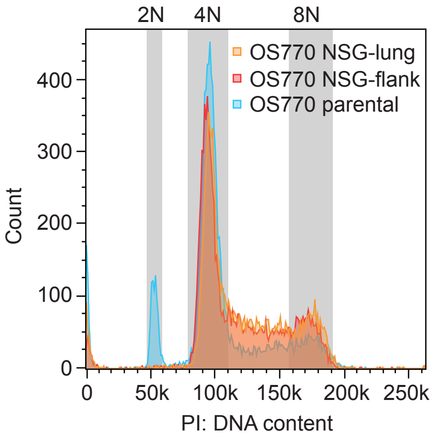


**F**

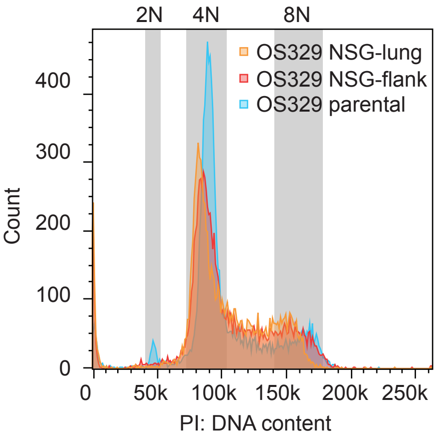

**Supplementary Figure S5. Karyotypic stability of OS cells across subcutaneous engraftment and lung metastatic colonization.**

(A) Spectral karyotyping of OS770 cells derived from the flank tumor in the NSG host.

(B) Spectral karyotyping of OS770 cells derived from the lung tumor in the NSG host.

(C) Spectral karyotyping of OS329 cells derived from the flank tumor in the NSG host.

(D) Spectral karyotyping of OS329 cells derived from the lung tumor in the NSG host.

(E) DNA content of parental OS770, NSG tumor or lung-derived OS770 cells was quantified by PI staining and flow cytometry.

(F) DNA content of parental OS329, NSG tumor or lung-derived OS329 cells was quantified by PI staining and flow cytometry.

St. Jude - Confidential

### Supplementary Figure S6

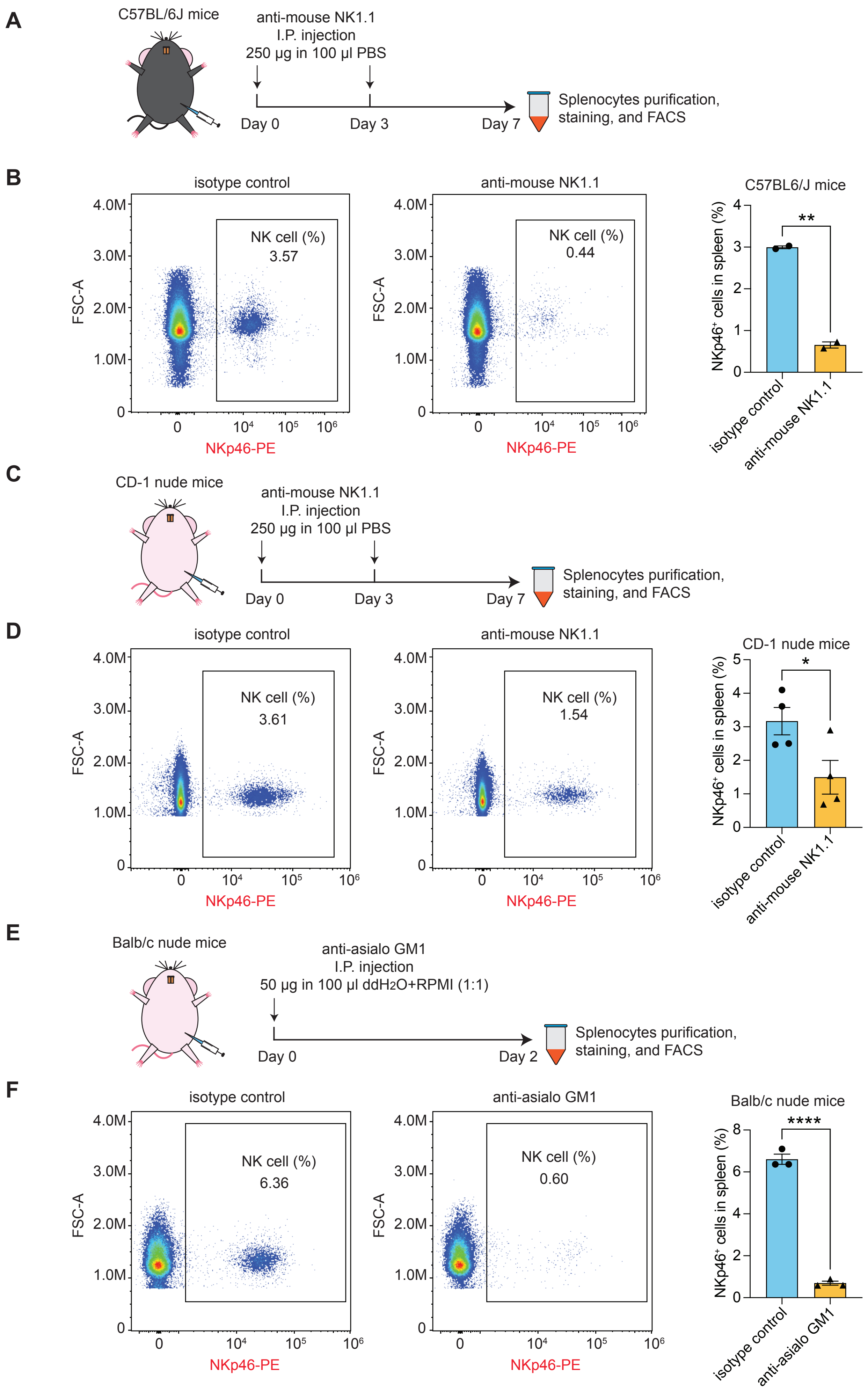

**Supplementary Figure S6. NK depletion efficacy in different mouse strains.**

(A) C57BL/6J mice were intraperitoneally (I.P.) injected with 250 µg of anti-mouse NK1.1 in 100 µL PBS or with the corresponding isotype control twice a week. Mice were euthanized on day 7 post-NK depletion, and cells were recovered from the spleen.

(B) Splenocytes were stained with anti-mouse NKp46 conjugated to phycoerythrin (PE). The percentage of NK cells in the spleen was assessed by flow cytometry, which showed a significant decrease following anti-mouse NK1.1 treatment in C57BL/6J mice.

(C) CD-1 nude mice were intraperitoneally injected with 250 µg of anti-mouse NK1.1 in 100 µL PBS or with the corresponding isotype control twice a week. Mice were euthanized on day 7 post-NK depletion, and the splenocytes were analyzed.

(D) The percentage of NK cells in the spleen was evaluated using the same strategy as in (B). Quantification showed a significantly reduced NK cell population in the spleens of CD-1 nude mice after anti-mouse NK1.1 treatment. However, the effects showed relatively large variability among individual mice.

(E) Balb/c nude mice were intraperitoneally injected with 50 µg anti-asialo GM1 in 100 µL of 50% dd$H_2$O and 50% RPMI. Mice were euthanized 2 days after NK depletion, and spleens were collected for splenocyte purification.

(F) Splenic NK cell population was determined by flow cytometry, which showed a dramatic decline of NK cells in the mice treated with anti-asialo GM1 compared to the isotype control.

Data were shown as means ± SDs; statistical analyses were performed using the Student's unpaired t-test: *$p < 0.05$, **$p < 0.01$, ***$p < 0.001$, ****$p < 0.0001$.

St. Jude - Confidential

### Supplementary Figure S7

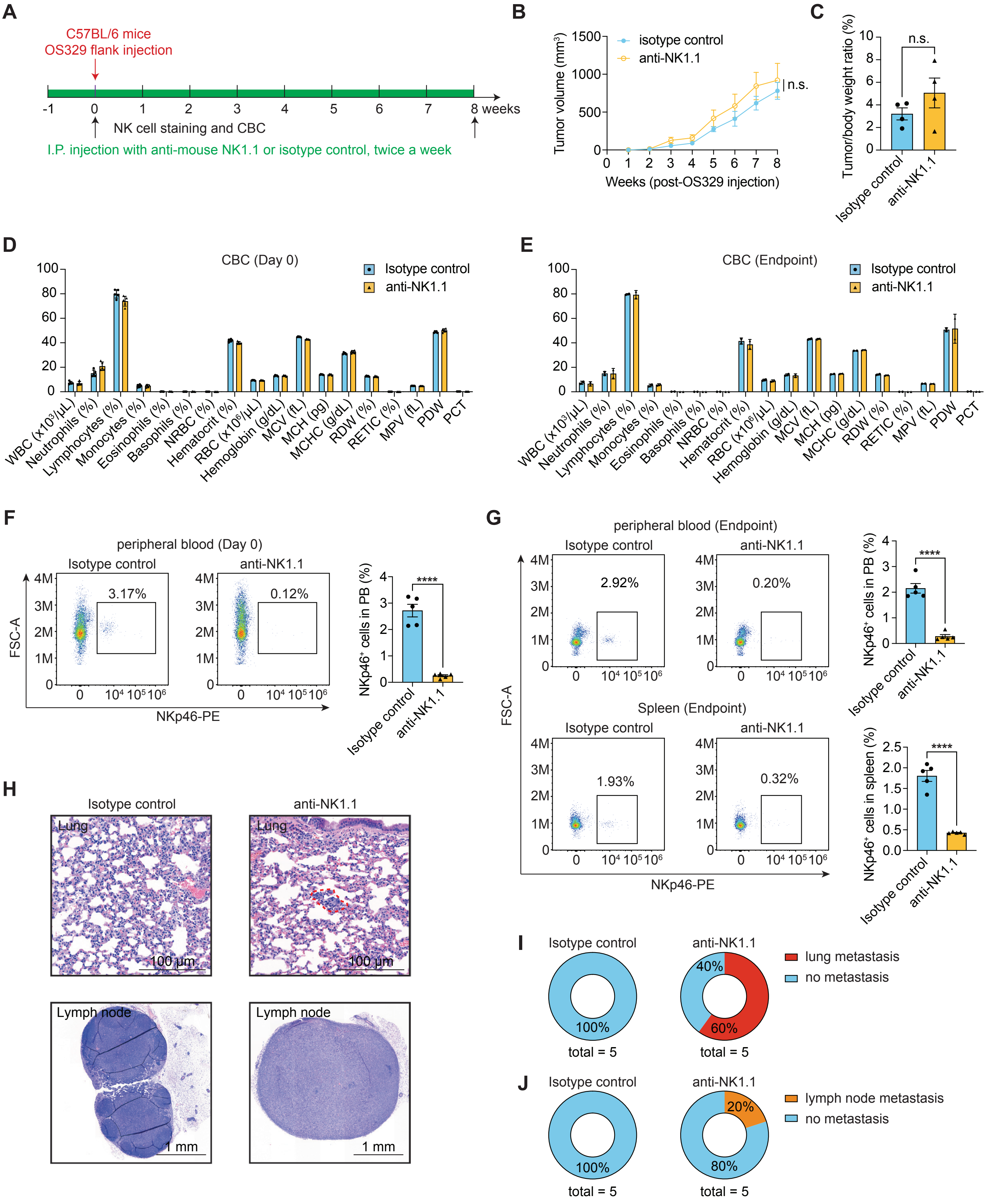

**Supplementary Figure S7. NK depletion promotes OS metastasis in C57BL/6J mice.**

(A) Regimen for NK depletion in C57BL/6J mice bearing flank OS tumors. Anti-mouse NK1.1 was injected intraperitoneally in five mice one week before OS329 implantation to neutralize NK cells. In the control group, five mice received IgG. Mice were then subcutaneously injected with $3 \times 10^6$ OS329 into the flank and treated with either 250 µg of anti-mouse NK1.1 or an isotype control twice a week. Tumor growth and lung metastasis were monitored weekly with a caliper. The efficacy of NK depletion in peripheral blood or spleen was assessed by flow cytometry at 1 week post-depletion and at the humane endpoint.

(B) Flank OS tumor growth curve.

(C) The ratio of tumor to body weight.

(D) Complete blood count (CBC) analysis for the peripheral blood (PB) collected via the submandibular vein one week after NK depletion.

(E) CBC for the PB collected via the submandibular vein at the humane endpoint.

(F) The efficacy of NK depletion was assessed by flow cytometry one week after NK depletion (day 0).

(G) The efficacy of NK depletion in the PB and spleen was assessed by flow cytometry at the humane endpoint.

(H) Representative histopathological sections of the lung and lymph node of the C57BL/6J mice bearing flank OS329 tumors that were treated with either isotype control or anti-mouse NK1.1.

(I) Pie charts showing the percentage of mice exhibiting lung metastasis.

(J) Pie charts showing the percentage of mice exhibiting lymph node metastasis.

In (B-G), data were shown as means ± SDs; statistical analyses were performed using the Student's unpaired t-test: $^{*}p < 0.05$, $^{***}p < 0.001$.

### Supplementary Figure S8

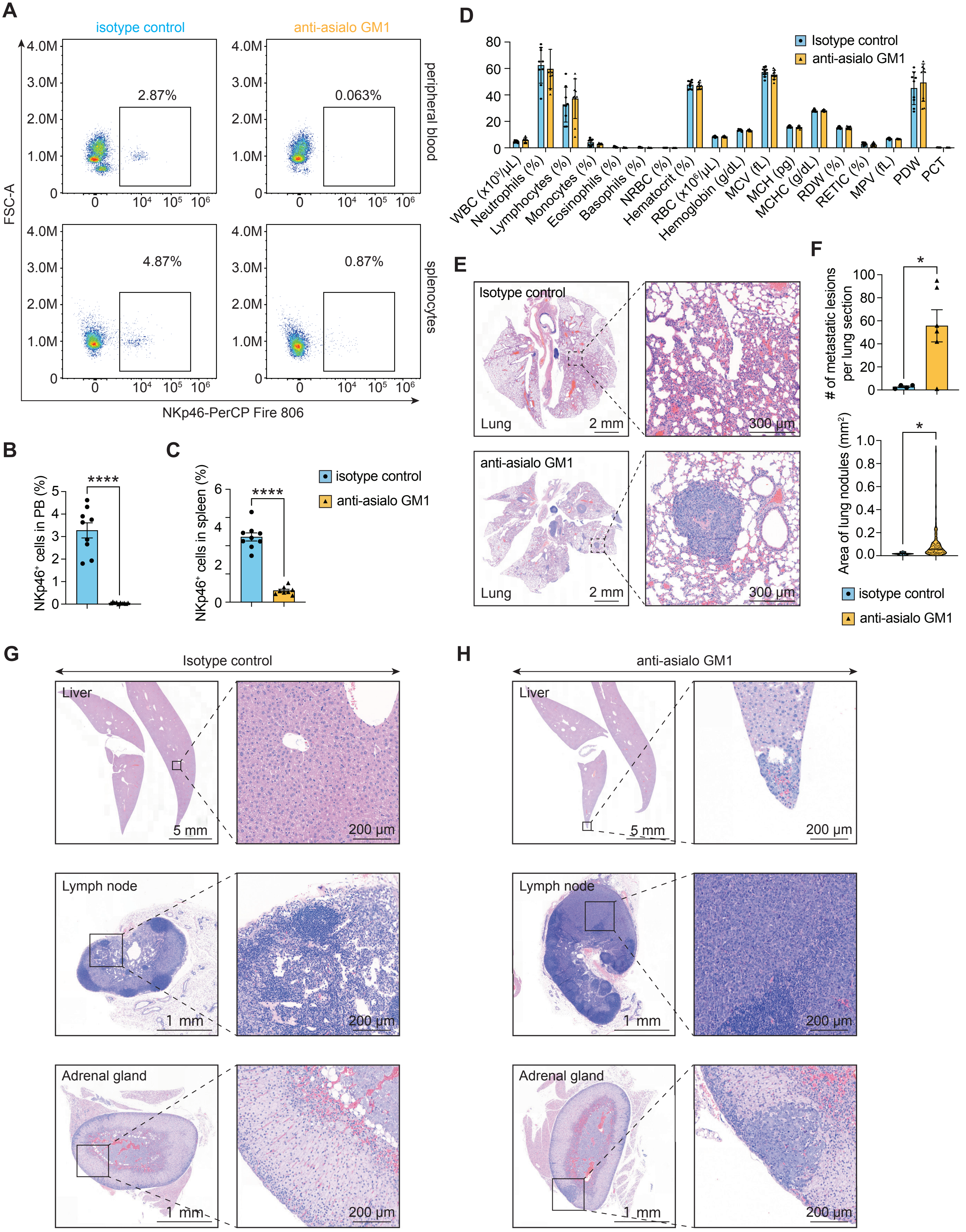

**Supplementary Figure S8. NK depletion promotes OS metastasis in Balb/c nude mice under a post-treatment regimen.**

After the establishment of luciferin-marked flank tumors, mice were treated with asialo GM1 antibody or an isotype control antibody and imaged until they reached the humane endpoint (See Figure 3G).

(A) The efficacy of NK depletion was assessed by flow cytometry at the humane endpoint.

(B) Quantification of the NK cell population in PB.

(C) Quantification of the NK cell population in the spleen.

(D) CBC of PB collected via the submandibular vein at the humane endpoint.

(E) Representative histopathological sections of the lung from Balb/c nude mice bearing flank OS tumors that were treated with either isotype control or anti-asialo GM1.

(F) Quantification of the number of lung lesions and the area of the lung metastatic nodules.

(G) Representative histopathological sections of liver, lymph node, and adrenal gland from Balb/c nude mice bearing flank OS tumors treated with an isotype control antibody.

(H) Representative histopathological sections from the Balb/c nude mice bearing flank OS tumors that were treated with an anti-asialo GM1 antibody.

Data from two independent animal cohorts were combined for statistical analysis: n (isotype control) = 4-9, n (anti-asialo GM1) = 5-10. In (B-D) and (F), data are shown as means ± SDs; statistical analyses were performed using the Student's unpaired t-test: *$p < 0.05$, ****$p < 0.0001$.

### Supplementary Figure S9

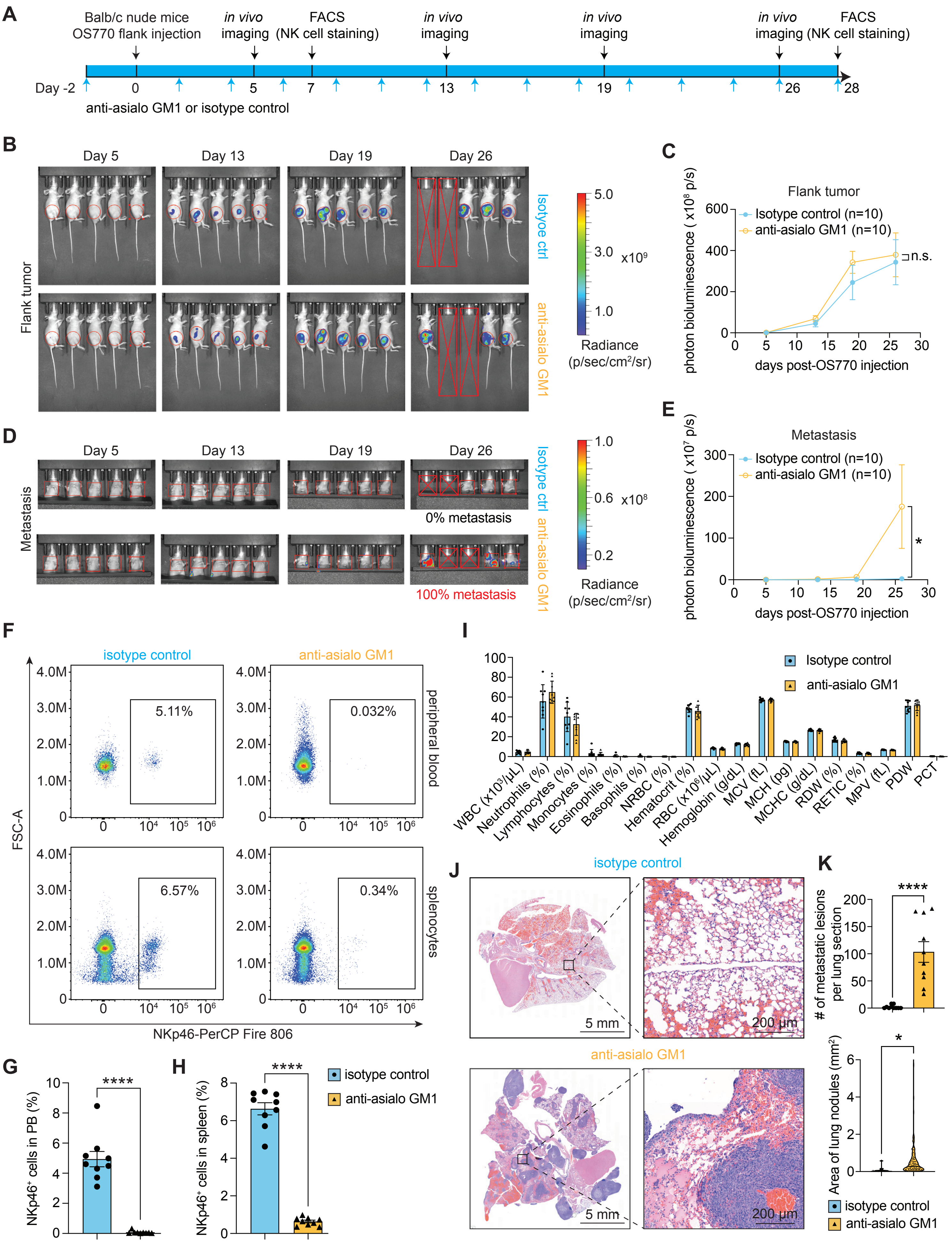

**Supplementary Figure S9. NK depletion promotes OS metastasis in Balb/c nude mice under a pre-treatment regimen.**

(A) Regimen for NK depletion in Balb/c nude mice bearing flank OS tumors. Anti-asialo GM1 was administered by intraperitoneal injection two days before OS implantation to neutralize NK cells. In the control group, mice received IgG. Mice were then subcutaneously injected with $3 \times 10^6$ OS770 into the flank and treated with either anti-asialo GM1 or isotype control every other day. Tumor growth and lung metastasis were monitored by bioluminescence imaging on days 5, 13, 19, and 26. The efficacy of NK depletion in peripheral blood or spleen was assessed by flow cytometry at one week after OS implantation and at the humane endpoint.

(B) Representative bioluminescence images of Balb/c nude mice bearing flank OS tumors.

(C) Quantification of signal intensity in flank OS tumors.

(D) Representative flank tumor-shield bioluminescence images of lung metastasis in Balb/c nude mice.

(E) Quantification of signal intensity of lung metastasis.

(F) The efficacy of NK depletion was assessed by flow cytometry at the humane endpoint.

(G) Quantification of the NK cell population in PB.

(H) Quantification of the NK cell population in the spleen.

(I) CBC for the PB collected via the submandibular vein at the humane endpoint.

(J) Representative histopathological sections of the lung of the Balb/c nude mice bearing flank OS tumors that were treated with either isotype control or anti-asialo GM1.

(K) Quantification of the number and area of lung metastatic lesions per section.

Data from two independent animal cohorts were combined for statistical analysis: n (isotype control) = 10, n (anti-asialo GM1) = 10. In (C), (E), (G-I), and (K), data are shown as means ± SDs; statistical analyses were performed using the Student's unpaired t-test: n.s. $p > 0.05$, *$p < 0.05$, ****$p < 0.0001$.

### Supplementary Figure S10

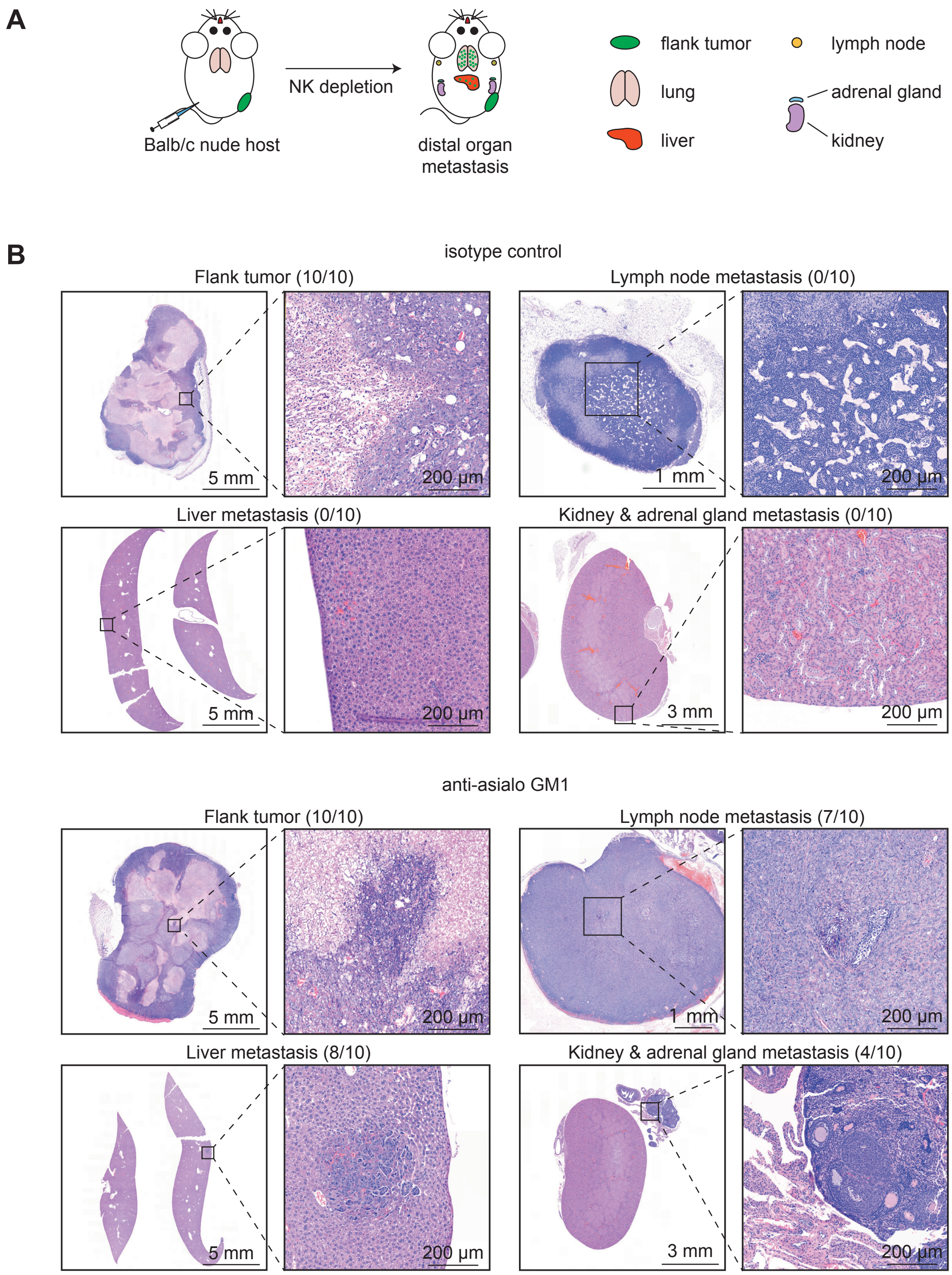

**Supplementary Figure S10. NK depletion promotes OS metastasis to distal organs in Balb/c nude mice bearing flank tumors.**

(A) Schematic of NK depletion facilitating OS metastasis to the distal organs, including the lung, liver, lymph node, adrenal gland, and kidney, in Balb/c nude mice injected with OS770 cells (described in **Supplementary Figure S9A**).

(B) Representative histopathological sections demonstrating that, compared with the isotype control group, NK cell depletion promoted OS tumor cell metastasis in distal organs.

St. Jude - Confidential

### Supplementary Figure S11

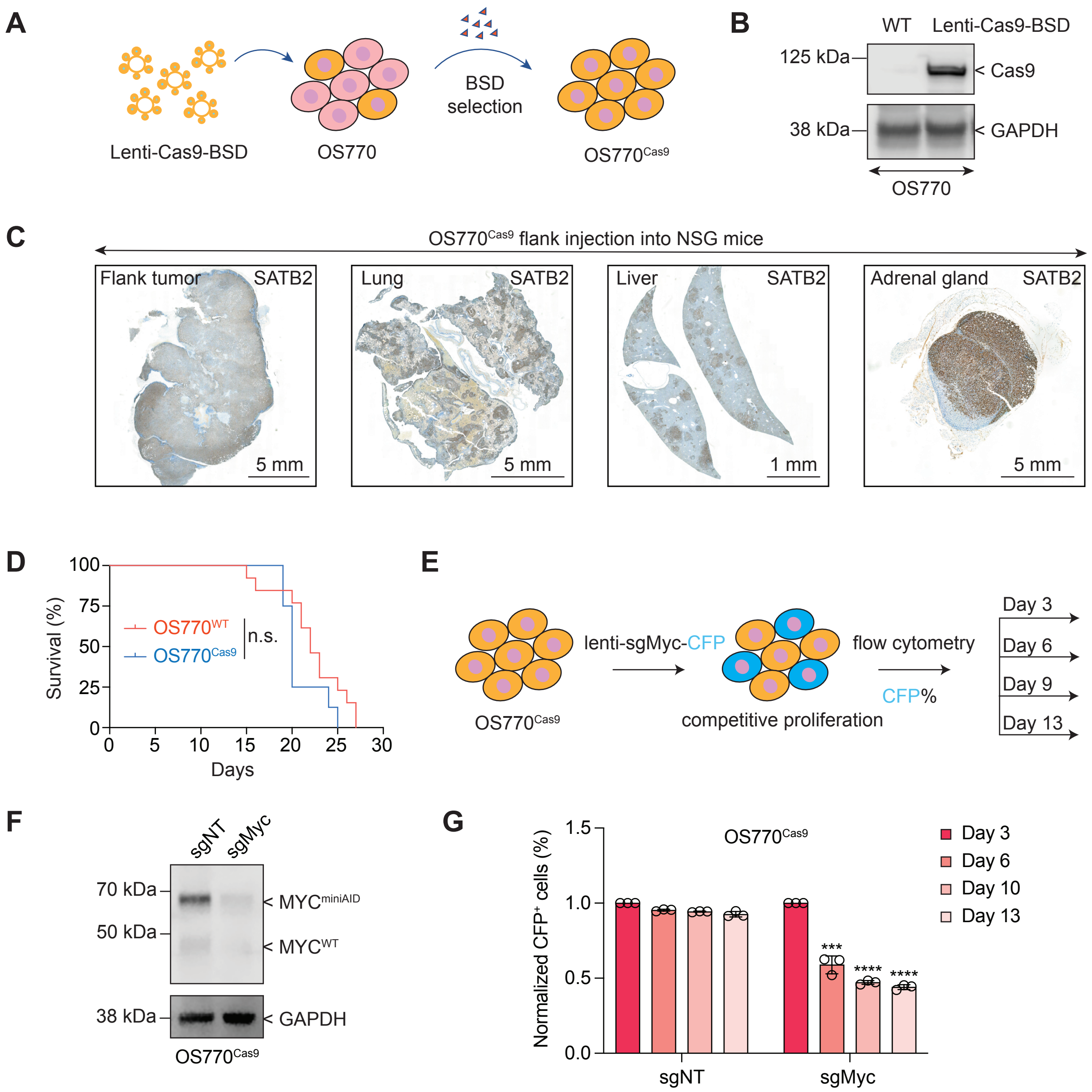

**Supplementary Figure S11. CRISPR/Cas9 editing of Myc in OS770 cells.**

(A) Schematic diagram of establishing the stably Cas9-expressing OS770 ($OS770^{Cas9}$) cell line. OS770 cells were transduced with a lentiviral vector for ectopic Cas9 expression, and stable Cas9-expressing cells were selected using blasticidin (BSD) based on the co-expressed BSD resistance gene.

(B) Immunoblots showed stable Cas9 expression in $OS770^{Cas9}$ cells.

(C) $OS770^{Cas9}$ cells were injected subcutaneously into the flank of NSG mice. Mice developed consistent bony OS flank tumors and distal organ metastasis revealed by SATB2 staining.

(D) Survival curves for NSG mice carrying $OS770^{WT}$ or $OS770^{Cas9}$ in the flank. Statistical analysis was performed using the log-rank test.

(E) Schematic of competitive proliferation assay (CPA). $OS770^{Cas9}$ cells were infected with lentiviruses encoding sgMyc with CFP fluorescence. The percentage of CFP-positive cells was traced by flow cytometry on days 3, 6, 9, and 13 post-infection.

(F) Immunoblotting showed that sgMyc effectively ablates Myc expression despite the presence of extra Myc copies in $OS770^{Cas9}$ cells.

(G) Competitive proliferation assay (CPA) showed that Myc knockout significantly impaired the cell fitness of OS770. Data were shown as means ± SDs; statistical analyses were performed using the Student’s unpaired t-test: ***$p$ < 0.001, ****$p$ < 0.0001.

### Supplementary Figure S12

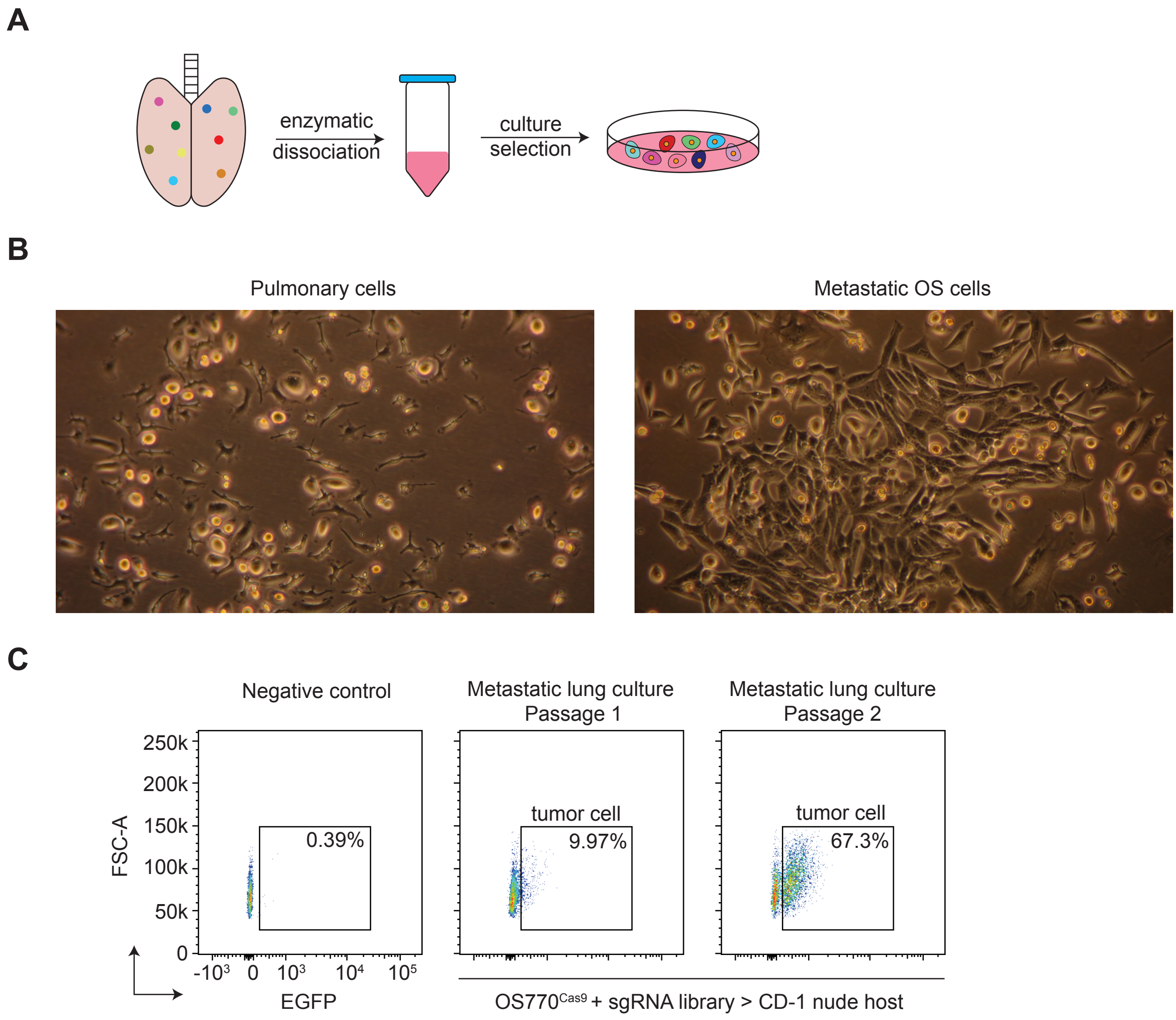

**Supplementary Figure S12. Optimized *ex vivo* culture conditions enrich metastatic OS cells from the lung.**

(A) Schematic diagram of the enrichment of pulmonary metastatic OS cells under *ex vivo* culture conditions. Metastatic lung tissue was harvested, enzymatically dissociated into a single-cell suspension, and cultured to selectively expand metastatic tumor cells.

(B) Representative image of cultured pulmonary cells (left; passage 1) and visual outgrowth of metastatic OS cells (right; passage 1).

(C) The percentage of metastatic OS cells in the culture condition was detected by flow cytometry.

St. Jude - Confidential

### Supplementary Figure S13

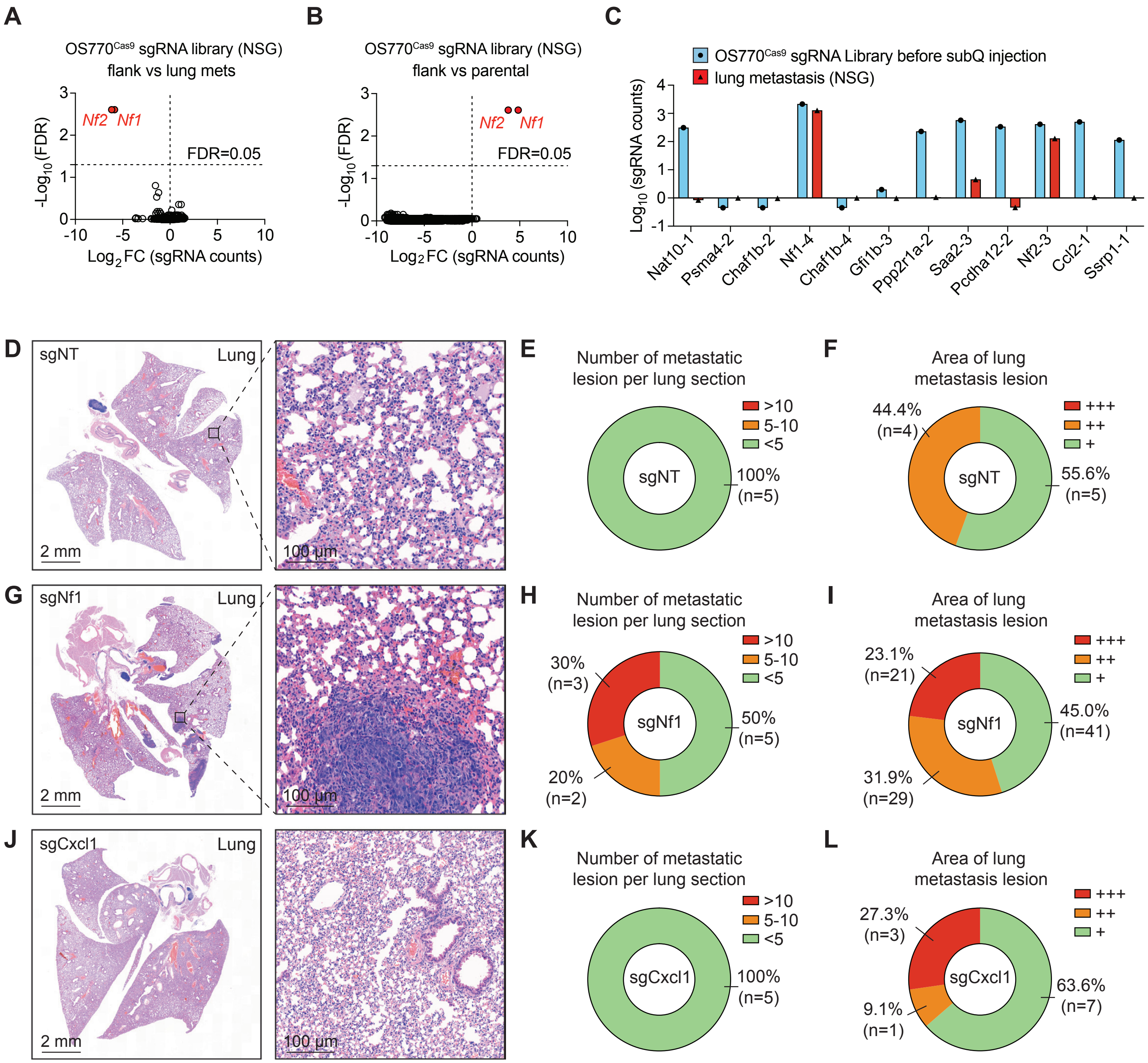

**Supplementary Figure S13. Whole-genome CRISPR screening identified that tumor suppressor *Nf1* controls OS lung metastasis.**

(A) Volcano plot of enriched sgRNAs in the lung of NSG mice compared to the flank tumor. The candidate genes that passed the statistical cutoff were highlighted.

(B) Volcano plot of enriched sgRNAs in the flank tumor of NSG mice compared to pooled OS770 cells before injection. The candidate genes that passed the statistical cutoff were highlighted.

(C) MAGeCK analysis was conducted in sgRNA library-induced lung metastatic cells from NSG mice compared to flank tumor cells.

(D) Representative histopathological H&E-stained lung sections from the CD-1 nude mice bearing subcutaneous OS770 flank tumors expressing sgNT.

(E) Pie charts showing the percentage of lung metastatic nodule numbers in each CD-1 nude mouse bearing subcutaneous OS770 flank tumors expressing sgNT. <5 metastatic nodules (green); 5-10 metastatic nodules (orange); >10 metastatic nodules (red).

(F) Pie charts showing the percentage of lung metastatic nodule areas in each CD-1 nude mouse bearing subcutaneous OS770 flank tumors expressing sgNT. + (green) area $\leq 0.01$ mm$^2$; ++ (orange) $0.01 <$ area $\leq 0.05$ mm$^2$; +++ (red) area $> 0.05$ mm$^2$.

(G) Representative histopathological H&E-stained lung sections from the CD-1 nude mice bearing subcutaneous OS770 flank tumors expressing sgNf1.

(H) Pie charts showing the percentage of lung metastatic nodule numbers in each CD-1 nude mouse bearing subcutaneous OS770 flank tumors expressing sgNf1. <5 metastatic nodules (green); 5-10 metastatic nodules (orange); >10 metastatic nodules (red).

(I) Pie charts showing the percentage of lung metastatic nodule areas in each CD-1 nude mouse bearing subcutaneous OS770 flank tumors expressing sgNf1. + (green) area $\leq 0.01$ mm$^2$; ++ (orange) $0.01 <$ area $\leq 0.05$ mm$^2$; +++ (red) area $> 0.05$ mm$^2$.

(J) Representative histopathological H&E-stained lung sections from the CD-1 nude mice bearing subcutaneous OS770 flank tumors expressing sgCxcl1.

(K) Pie charts showing the percentage of lung metastatic nodule numbers in each CD-1 nude mouse bearing subcutaneous OS770 flank tumors expressing sgCxcl1. <5 metastatic nodules (green); 5-10 metastatic nodules (orange); >10 metastatic nodules (red).

(L) Pie charts showing the percentage of lung metastatic nodule areas in each CD-1 nude mouse bearing subcutaneous OS770 flank tumors expressing sgCxcl1. + (green) area $\leq 0.01$ mm$^2$; ++ (orange) $0.01 <$ area $\leq 0.05$ mm$^2$; +++ (red) area $> 0.05$ mm$^2$.

### Supplementary Figure S14

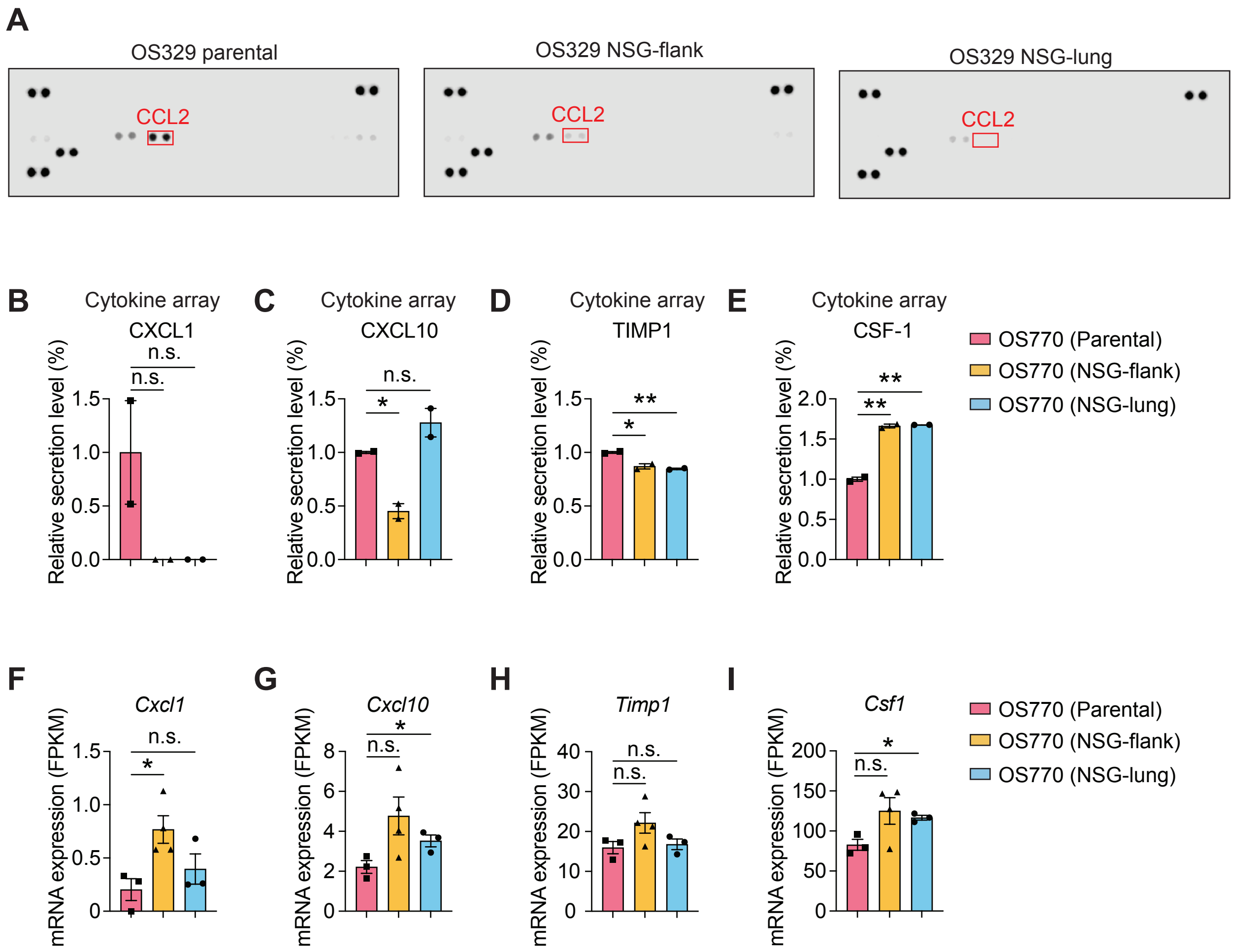

**Supplementary Figure S14. OS tumor cells recovered from NSG hosts showed decreased CCL2 at both the mRNA expression and protein secretion levels.**

(A) Cytokine arrays for parental OS329 cells, NSG flank tumor-derived OS329 cells, and NSG lung tumor-derived OS329 cells.

Cytokine arrays for parental OS770 cells, NSG flank tumor-derived OS770 cells, and NSG lung tumor-derived OS770 cells are shown in Figure 5D, 5F, and 5G. The quantification of each secreted factor is listed below:

(B) The quantification of secreted CXCL1.

(C) The quantification of secreted CXCL10.

(D) The quantification of secreted TIMP1.

(E) The quantification of secreted CSF-1.

(F) *Cxcl1* mRNA expression levels were detected by RNA-seq and normalized using FPKM.

(G) *Cxcl10* mRNA expression levels were detected by RNA-seq and normalized using FPKM.

(H) *Timp1* mRNA expression levels were detected by RNA-seq and normalized using FPKM.

(I) *Csf1* mRNA expression levels were detected by RNA-seq and normalized using FPKM.

In (B-I), data were shown as means ± SDs; statistical analyses were performed using the Student's unpaired t-test: n.s. $p > 0.05$, *$p < 0.05$, **$p < 0.01$.

St. Jude - Confidential

### Supplementary Figure S15

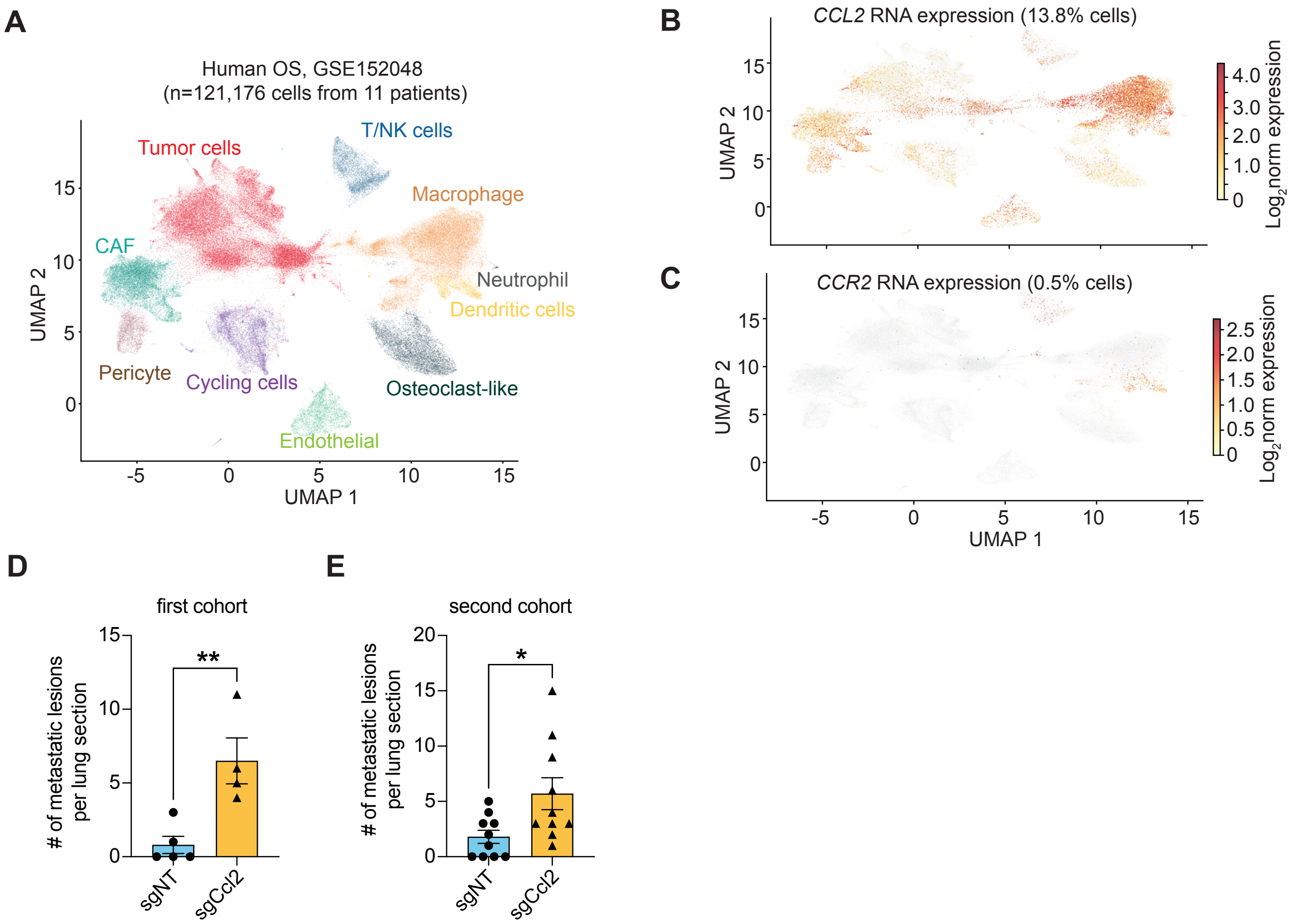

**Supplementary Figure S15. CCL2-CCR2 signaling within OS TME.**

(A) Single-cell RNA-seq data collected from 11 OS patients were downloaded from GSE152048 and re-analyzed. A total of 121,176 passed quality control and were used to annotate distinct cell populations based on marker gene expression, visualized using uniform manifold approximation and projection (UMAP).

(B) The expression of *CCL2* across different cell types.

(C) The expression of *CCR2* across different cell types.

(D) Quantification of the number of pulmonary metastatic nodules in the first CD-1 nude mice injected with sgNT (n = 5) or sgCcl2 (n = 4) OS770 cells cohort.

(E) Quantification of the number of pulmonary metastatic nodules in the second CD-1 nude mice injected with sgNT (n = 10) or sgCcl2 (n = 10) OS770 cells cohort.

In (D) and (E), data were shown as means ± SDs; statistical analyses were performed using the Student's unpaired t-test: n.s. $p > 0.05$, *$p < 0.05$, **$p < 0.01$.

**Supplementary Table S1. Summary of tumor phenotypes generated by *Trp53*-null, MYC-overexpressing sphere cells.**

Summary of 33 primary tumors arising in CD-1 nude recipient mice injected with tumor-initiating sphere cells carrying *Trp53*-null alleles combined with either Lenti-Myc-RFP or MSCV-Myc-HA-miniAID-IRES-OsTIR1$^{F74G}$-P2A-EGFP transgenes into the cortices. For each animal, the table lists the mouse ID, engineered genotype, tumor type (OS, osteosarcoma; MB, medulloblastoma; undefined), survival in days from tumor onset to the humane endpoint, and immunohistochemical status of SATB2 (osteoblast lineage marker) and PTEN (tumor suppressor).

| Mouse ID | Engineered Genotype | Tumor Types | Survival (days) | SATB2 | PTEN |
|---|---|---|---|---|---|
| 322 | *Trp53*$^{-/-}$; Lenti-Myc-RFP | MB | 84 | negative | positive |
| 773 | *Trp53*$^{-/-}$; MSCV-Myc-HA-miniAID-IRES-OsTIR1$^{F74G}$-P2A-EGFP | MB | 90 | negative | positive |
| 784 | *Trp53*$^{-/-}$; MSCV-Myc-HA-miniAID-IRES-OsTIR1$^{F74G}$-P2A-EGFP | MB | 111 | negative | positive |
| 73878 | *Trp53*$^{-/-}$; Lenti-Myc-RFP | MB | 34 | negative | positive |
| 73882 | *Trp53*$^{-/-}$; Lenti-Myc-RFP | MB | 59 | negative | positive |
| 73885 | *Trp53*$^{-/-}$; Lenti-Myc-RFP | MB | 91 | negative | positive |
| 323 | *Trp53*$^{-/-}$; Lenti-Myc-RFP | OS | 113 | positive | negative |
| 327 | *Trp53*$^{-/-}$; Lenti-Myc-RFP | OS | 302 | positive | negative |
| 329 | *Trp53*$^{-/-}$; Lenti-Myc-RFP | OS | 195 | positive | positive |

| | | | | | |
|---|---|---|---|---|---|
| 330 | *Trp53*$^{-/-}$; Lenti-Myc-RFP | OS | 197 | positive | positive |
| 765 | *Trp53*$^{-/-}$; MSCV-Myc-HA-miniAID-IRES-OsTIR1$^{F74G}$-P2A-EGFP | OS | 91 | positive | positive |
| 767 | *Trp53*$^{-/-}$; MSCV-Myc-HA-miniAID-IRES-OsTIR1$^{F74G}$-P2A-EGFP | OS | 106 | positive | positive |
| 768 | *Trp53*$^{-/-}$; MSCV-Myc-HA-miniAID-IRES-OsTIR1$^{F74G}$-P2A-EGFP | OS | 186 | positive | negative |
| 769 | *Trp53*$^{-/-}$; MSCV-Myc-HA-miniAID-IRES-OsTIR1$^{F74G}$-P2A-EGFP | OS | 151 | positive | positive |
| 770 | *Trp53*$^{-/-}$; MSCV-Myc-HA-miniAID-IRES-OsTIR1$^{F74G}$-P2A-EGFP | OS | 153 | positive | negative |
| 771 | *Trp53*$^{-/-}$; MSCV-Myc-HA-miniAID-IRES-OsTIR1$^{F74G}$-P2A-EGFP | OS | 208 | positive | negative |
| 772 | *Trp53*$^{-/-}$; MSCV-Myc-HA-miniAID-IRES-OsTIR1$^{F74G}$-P2A-EGFP | OS | 96 | positive | positive |
| 774 | *Trp53*$^{-/-}$; MSCV-Myc-HA-miniAID- | OS | 181 | positive | negative |

| | | | | | |
|---|---|---|---|---|---|
| | IRES-OsTIR1$^{F74G}$-P2A-EGFP | | | | |
| 775 | *Trp53*$^{-/-}$; MSCV-Myc-HA-miniAID-IRES-OsTIR1$^{F74G}$-P2A-EGFP | OS | 119 | positive | positive |
| 776 | *Trp53*$^{-/-}$; MSCV-Myc-HA-miniAID-IRES-OsTIR1$^{F74G}$-P2A-EGFP | OS | 82 | positive | low expression |
| 777 | *Trp53*$^{-/-}$; MSCV-Myc-HA-miniAID-IRES-OsTIR1$^{F74G}$-P2A-EGFP | OS | 105 | positive | positive |
| 779 | *Trp53*$^{-/-}$; MSCV-Myc-HA-miniAID-IRES-OsTIR1$^{F74G}$-P2A-EGFP | OS | 89 | positive | positive |
| 780 | *Trp53*$^{-/-}$; MSCV-Myc-HA-miniAID-IRES-OsTIR1$^{F74G}$-P2A-EGFP | OS | 175 | positive | negative |
| 781 | *Trp53*$^{-/-}$; MSCV-Myc-HA-miniAID-IRES-OsTIR1$^{F74G}$-P2A-EGFP | OS | 97 | positive | negative |
| 782 | *Trp53*$^{-/-}$; MSCV-Myc-HA-miniAID-IRES-OsTIR1$^{F74G}$-P2A-EGFP | OS | 232 | NA | NA |
| 783 | *Trp53*$^{-/-}$; MSCV-Myc-HA-miniAID- | OS | 54 | positive | low expression |

| | | | | | |
|---|---|---|---|---|---|
| | IRES-OsTIR1$^{F74G}$-P2A-EGFP | | | | |
| 73879 | *Trp53*$^{-/-}$; Lenti-Myc-RFP | OS | 148 | positive | positive |
| 73880 | *Trp53*$^{-/-}$; Lenti-Myc-RFP | OS | 142 | positive | positive |
| 73883 | *Trp53*$^{-/-}$; Lenti-Myc-RFP | OS | 93 | positive | low expression |
| 73884 | *Trp53*$^{-/-}$; Lenti-Myc-RFP | OS | 161 | positive | positive |
| 73887 | *Trp53*$^{-/-}$; Lenti-Myc-RFP | OS | 92 | positive | positive |
| 73881 | *Trp53*$^{-/-}$; Lenti-Myc-RFP | undefined | 92 | - | - |
| 73886 | *Trp53*$^{-/-}$; Lenti-Myc-RFP | undefined | 91 | - | - |

**Supplementary Table S2. Genome-wide ectopic *Myc* integration sites identified by whole-genome sequencing (WGS).**

Integration site analysis of Myc-encoding vectors across eight samples, including the tumor-initiating sphere cell, parental OS770 cells, OS770 cells derived from the flank tumor of NSG host (OS770-NSG-flank), OS770 cells derived from the lung tumor of NSG host (OS770-NSG-lung), parental OS329 cells, OS329 cells derived from the flank tumor of NSG host (OS329-NSG-flank), OS329 cells derived from the lung tumor of NSG host (OS329-NSG-lung) and OS329 cells derived from the flank tumor of C57BL/6J host (OS329-BL6-flank). Each sheet corresponds to one sample. Integration clusters were identified from reads in WGS data and are reported with the following fields: Cluster_id (cluster identifier, each cluster represents an integration site), CHROM/START/END (genomic coordinates, mm10), and READ (total read count).

| **Sphere cells + Myc (tumor-initiating cells)** | | | | |
|---|---|---|---|---|
| **Cluster_id** | **CHROM** | **START** | **END** | **READ** |
| Cluster_2 | chr2 | 26506959 | 26507165 | 14 |
| Cluster_8 | chr15 | 73455841 | 73455905 | 4 |
| Cluster_10 | chr19 | 5464811 | 5464910 | 3 |
| Cluster_7 | chr14 | 105739599 | 105739718 | 3 |
| Cluster_1 | chr1 | 88544201 | 88544266 | 2 |
| Cluster_5 | chr11 | 49445334 | 49445400 | 2 |
| Cluster_6 | chr11 | 98710223 | 98710274 | 2 |
| Cluster_3 | chr3 | 127298670 | 127298710 | 1 |
| Cluster_4 | chr11 | 6006358 | 6006406 | 1 |
| Cluster_9 | chr16 | 91255714 | 91255830 | 1 |
| **OS770-parental** | | | | |
| **Cluster_id** | **CHROM** | **START** | **END** | **READ** |
| Cluster_12 | chr12 | 29873029 | 29873247 | 50 |
| Cluster_11 | chr11 | 94933101 | 94933320 | 33 |
| Cluster_9 | chr11 | 6006298 | 6006408 | 21 |
| Cluster_18 | chr18 | 66488938 | 66488978 | 5 |
| Cluster_8 | chr10 | 19595628 | 19595658 | 5 |
| Cluster_16 | chr17 | 87106120 | 87106161 | 4 |
| Cluster_10 | chr11 | 6011425 | 6011514 | 3 |
| Cluster_14 | chr14 | 19729487 | 19729555 | 2 |
| Cluster_15 | chr14 | 25832967 | 25833037 | 2 |
| Cluster_17 | chr18 | 52639319 | 52639423 | 2 |
| Cluster_7 | chr8 | 91287824 | 91287854 | 2 |
| Cluster_1 | chr2 | 30919003 | 30919102 | 1 |

| | | | | |
|---|---|---|---|---|
| Cluster_13 | chr12 | 80228554 | 80228673 | 1 |
| Cluster_19 | chrX | 57305762 | 57305802 | 1 |
| Cluster_2 | chr3 | 49641790 | 49641849 | 1 |
| Cluster_20 | chrX | 103494839 | 103494902 | 1 |
| Cluster_3 | chr4 | 126154974 | 126155042 | 1 |
| Cluster_4 | chr5 | 53489671 | 53489751 | 1 |
| Cluster_5 | chr5 | 120551939 | 120552016 | 1 |
| Cluster_6 | chr6 | 4491356 | 4491434 | 1 |
| | | **OS770-NSG-flank** | | |
| **Cluster_id** | **CHROM** | **START** | **END** | **READ** |
| Cluster_21 | chr11 | 94933102 | 94933337 | 45 |
| Cluster_23 | chr12 | 29873022 | 29873260 | 41 |
| Cluster_28 | chr15 | 10815422 | 10815603 | 16 |
| Cluster_33 | chr19 | 10267119 | 10267194 | 9 |
| Cluster_16 | chr9 | 50750562 | 50750596 | 6 |
| Cluster_7 | chr4 | 64201334 | 64201409 | 6 |
| Cluster_1 | chr1 | 55900550 | 55900712 | 4 |
| Cluster_8 | chr4 | 89427729 | 89427768 | 4 |
| Cluster_13 | chr8 | 87492397 | 87492561 | 3 |
| Cluster_17 | chr10 | 8385091 | 8385208 | 3 |
| Cluster_26 | chr12 | 108304159 | 108304334 | 3 |
| Cluster_3 | chr2 | 30743577 | 30743644 | 3 |
| Cluster_30 | chr15 | 62360547 | 62360720 | 3 |
| Cluster_19 | chr11 | 6006340 | 6006406 | 2 |
| Cluster_2 | chr1 | 90839423 | 90839539 | 2 |
| Cluster_22 | chr11 | 98748263 | 98748377 | 2 |
| Cluster_27 | chr13 | 23582423 | 23582526 | 2 |
| Cluster_31 | chr16 | 86035233 | 86035334 | 2 |
| Cluster_32 | chr16 | 95533406 | 95533489 | 2 |
| Cluster_5 | chr3 | 95985749 | 95985817 | 2 |
| Cluster_6 | chr4 | 64085728 | 64085848 | 2 |
| Cluster_9 | chr5 | 21785940 | 21786008 | 2 |
| Cluster_10 | chr5 | 137106040 | 137106071 | 1 |
| Cluster_11 | chr8 | 61796843 | 61796878 | 1 |
| Cluster_12 | chr8 | 82953337 | 82953381 | 1 |
| Cluster_14 | chr8 | 123426120 | 123426157 | 1 |
| Cluster_15 | chr8 | 127317983 | 127318078 | 1 |
| Cluster_18 | chr10 | 81176266 | 81176367 | 1 |
| Cluster_20 | chr11 | 6011444 | 6011514 | 1 |
| Cluster_24 | chr12 | 72748903 | 72748959 | 1 |
| Cluster_25 | chr12 | 104983221 | 104983302 | 1 |

| | | | | |
|---|---|---|---|---|
| Cluster_29 | chr15 | 38202908 | 38203020 | 1 |
| Cluster_34 | chrX | 36748405 | 36748437 | 1 |
| Cluster_4 | chr3 | 88617311 | 88617397 | 1 |
| **OS770-NSG-lung** | | | | |
| **Cluster_id** | **CHROM** | **START** | **END** | **READ** |
| Cluster_19 | chr11 | 94933107 | 94933315 | 44 |
| Cluster_22 | chr12 | 29873075 | 29873239 | 30 |
| Cluster_16 | chr11 | 6006292 | 6006406 | 13 |
| Cluster_18 | chr11 | 70667566 | 70667771 | 7 |
| Cluster_28 | chr16 | 86035201 | 86035333 | 6 |
| Cluster_33 | chrX | 142897976 | 142898133 | 6 |
| Cluster_29 | chr16 | 95533406 | 95533517 | 5 |
| Cluster_23 | chr12 | 108304247 | 108304325 | 4 |
| Cluster_1 | chr1 | 55900586 | 55900753 | 3 |
| Cluster_10 | chr6 | 129229688 | 129229804 | 3 |
| Cluster_14 | chr8 | 87492469 | 87492581 | 3 |
| Cluster_17 | chr11 | 6011438 | 6011514 | 3 |
| Cluster_26 | chr16 | 4881380 | 4881425 | 3 |
| Cluster_3 | chr4 | 21555529 | 21555626 | 3 |
| Cluster_6 | chr4 | 89427633 | 89427731 | 3 |
| Cluster_8 | chr5 | 21785948 | 21786008 | 3 |
| Cluster_2 | chr2 | 30743486 | 30743581 | 2 |
| Cluster_31 | chrX | 36748433 | 36748483 | 2 |
| Cluster_11 | chr7 | 110003031 | 110003091 | 1 |
| Cluster_12 | chr7 | 133136700 | 133136745 | 1 |
| Cluster_13 | chr8 | 23567668 | 23567699 | 1 |
| Cluster_15 | chr9 | 7267719 | 7267765 | 1 |
| Cluster_20 | chr11 | 96822387 | 96822443 | 1 |
| Cluster_21 | chr11 | 98748337 | 98748377 | 1 |
| Cluster_24 | chr13 | 23582416 | 23582526 | 1 |
| Cluster_25 | chr15 | 10815391 | 10815498 | 1 |
| Cluster_27 | chr16 | 64872691 | 64872752 | 1 |
| Cluster_30 | chr19 | 10267119 | 10267225 | 1 |
| Cluster_32 | chrX | 48083745 | 48083859 | 1 |
| Cluster_4 | chr4 | 55730811 | 55730911 | 1 |
| Cluster_5 | chr4 | 64085728 | 64085787 | 1 |
| Cluster_7 | chr4 | 152189731 | 152189801 | 1 |
| Cluster_9 | chr6 | 87006581 | 87006661 | 1 |
| **OS329-parental** | | | | |
| **Cluster_id** | **CHROM** | **START** | **END** | **READ** |
| Cluster_17 | chr14 | 122365064 | 122365282 | 51 |

| | | | | |
|---|---|---|---|---|
| Cluster_15 | chr11 | 4794109 | 4794335 | 47 |
| Cluster_12 | chr10 | 40482275 | 40482500 | 39 |
| Cluster_4 | chr6 | 4927215 | 4927441 | 39 |
| Cluster_16 | chr11 | 119792509 | 119792733 | 35 |
| Cluster_13 | chr10 | 99164007 | 99164238 | 30 |
| Cluster_19 | chr18 | 24512466 | 24512687 | 30 |
| Cluster_2 | chr2 | 35319554 | 35319772 | 30 |
| Cluster_9 | chr8 | 12820034 | 12820264 | 30 |
| Cluster_11 | chr8 | 127464822 | 127465021 | 28 |
| Cluster_6 | chr7 | 19379221 | 19379437 | 24 |
| Cluster_14 | chr10 | 119207699 | 119207900 | 23 |
| Cluster_20 | chr18 | 73654404 | 73654599 | 22 |
| Cluster_7 | chr7 | 40634194 | 40634409 | 22 |
| Cluster_8 | chr7 | 134167947 | 134168171 | 21 |
| Cluster_10 | chr8 | 113051997 | 113052207 | 20 |
| Cluster_21 | chr18 | 85684882 | 85685080 | 13 |
| Cluster_18 | chr18 | 19956681 | 19956897 | 11 |
| Cluster_22 | chrX | 123453274 | 123453395 | 8 |
| Cluster_1 | chr1 | 130514963 | 130515019 | 1 |
| Cluster_23 | chrX | 123894901 | 123894951 | 1 |
| Cluster_3 | chr4 | 124844037 | 124844376 | 1 |
| Cluster_5 | chr6 | 100928285 | 100928409 | 1 |
| **OS329-NSG-flank** | | | | |
| **Cluster_id** | **CHROM** | **START** | **END** | **READ** |
| Cluster_4 | chr7 | 19379212 | 19379433 | 34 |
| Cluster_11 | chr10 | 40482279 | 40482483 | 20 |
| Cluster_15 | chr11 | 119792511 | 119792724 | 19 |
| Cluster_20 | chr18 | 73654414 | 73654616 | 19 |
| Cluster_3 | chr6 | 4927221 | 4927430 | 19 |
| Cluster_16 | chr14 | 122365058 | 122365280 | 18 |
| Cluster_14 | chr11 | 4794104 | 4794338 | 17 |
| Cluster_6 | chr7 | 134167946 | 134168177 | 17 |
| Cluster_19 | chr18 | 24512468 | 24512675 | 16 |
| Cluster_13 | chr10 | 119207683 | 119207910 | 15 |
| Cluster_21 | chr18 | 85684866 | 85685095 | 15 |
| Cluster_8 | chr8 | 113052008 | 113052215 | 15 |
| Cluster_12 | chr10 | 99164023 | 99164233 | 14 |
| Cluster_7 | chr8 | 12820042 | 12820267 | 14 |
| Cluster_18 | chr18 | 19956686 | 19956903 | 13 |
| Cluster_5 | chr7 | 40634192 | 40634386 | 12 |
| Cluster_10 | chr8 | 127464884 | 127465030 | 7 |

| | | | | |
|---|---|---|---|---|
| Cluster_17 | chr15 | 11410286 | 11410395 | 5 |
| Cluster_22 | chr19 | 24673824 | 24673856 | 5 |
| Cluster_1 | chr2 | 35319574 | 35319669 | 4 |
| Cluster_25 | chrX | 123894901 | 123895009 | 4 |
| Cluster_9 | chr8 | 117128434 | 117128513 | 2 |
| Cluster_2 | chr3 | 148447761 | 148447875 | 1 |
| Cluster_23 | chr19 | 46984471 | 46984533 | 1 |
| Cluster_24 | chrX | 123453394 | 123453443 | 1 |
| | | **OS329-NSG-lung** | | |
| **Cluster_id** | **CHROM** | **START** | **END** | **READ** |
| Cluster_24 | chr18 | 24512464 | 24512692 | 58 |
| Cluster_9 | chr8 | 127464866 | 127465043 | 56 |
| Cluster_4 | chr7 | 19379211 | 19379439 | 53 |
| Cluster_13 | chr11 | 4794109 | 4794343 | 52 |
| Cluster_7 | chr8 | 12820041 | 12820262 | 47 |
| Cluster_6 | chr7 | 134167968 | 134168174 | 44 |
| Cluster_23 | chr18 | 19956673 | 19956894 | 40 |
| Cluster_1 | chr2 | 35319550 | 35319769 | 37 |
| Cluster_10 | chr10 | 40482284 | 40482500 | 37 |
| Cluster_5 | chr7 | 40634190 | 40634400 | 33 |
| Cluster_18 | chr14 | 122365057 | 122365266 | 30 |
| Cluster_12 | chr10 | 119207702 | 119207862 | 26 |
| Cluster_2 | chr6 | 4927216 | 4927411 | 18 |
| Cluster_11 | chr10 | 99164041 | 99164238 | 16 |
| Cluster_17 | chr13 | 88453032 | 88453536 | 16 |
| Cluster_8 | chr8 | 113052024 | 113052215 | 14 |
| Cluster_26 | chr18 | 73654392 | 73654599 | 13 |
| Cluster_16 | chr11 | 119792507 | 119792709 | 11 |
| Cluster_27 | chr18 | 85684878 | 85685090 | 11 |
| Cluster_28 | chrX | 123894901 | 123895007 | 10 |
| Cluster_14 | chr11 | 7834243 | 7834283 | 1 |
| Cluster_15 | chr11 | 70340920 | 70340962 | 1 |
| Cluster_19 | chr17 | 29588984 | 29589530 | 1 |
| Cluster_20 | chr17 | 47712764 | 47712832 | 1 |
| Cluster_21 | chr17 | 74289576 | 74289643 | 1 |
| Cluster_22 | chr18 | 7491319 | 7491383 | 1 |
| Cluster_25 | chr18 | 59206035 | 59206168 | 1 |
| Cluster_3 | chr6 | 135150695 | 135150768 | 1 |
| | | **OS329-BL6-flank** | | |
| **Cluster_id** | **CHROM** | **START** | **END** | **READ** |
| Cluster_12 | chr10 | 99164006 | 99164222 | 22 |

| | | | | |
|---|---|---|---|---|
| Cluster_2 | chr6 | 4927236 | 4927437 | 19 |
| Cluster_9 | chr8 | 127464872 | 127465024 | 19 |
| Cluster_7 | chr8 | 113051996 | 113052215 | 18 |
| Cluster_14 | chr11 | 4794118 | 4794312 | 15 |
| Cluster_3 | chr7 | 19379237 | 19379434 | 13 |
| Cluster_22 | chr18 | 73654234 | 73654598 | 10 |
| Cluster_13 | chr10 | 119207691 | 119207871 | 9 |
| Cluster_15 | chr11 | 119792536 | 119792682 | 9 |
| Cluster_20 | chr18 | 19956710 | 19956840 | 9 |
| Cluster_21 | chr18 | 24512472 | 24512675 | 9 |
| Cluster_5 | chr7 | 134167968 | 134168173 | 9 |
| Cluster_11 | chr10 | 40482302 | 40482477 | 7 |
| Cluster_17 | chr14 | 122365121 | 122365263 | 7 |
| Cluster_6 | chr8 | 12820085 | 12820228 | 7 |
| Cluster_23 | chr18 | 85684920 | 85685040 | 5 |
| Cluster_24 | chrX | 123894901 | 123894990 | 4 |
| Cluster_4 | chr7 | 40634240 | 40634307 | 3 |
| Cluster_1 | chr3 | 75052823 | 75263642 | 1 |
| Cluster_10 | chr9 | 63084601 | 63084704 | 1 |
| Cluster_16 | chr13 | 58017822 | 58017888 | 1 |
| Cluster_18 | chr17 | 33020545 | 33020603 | 1 |
| Cluster_19 | chr18 | 16488017 | 16488075 | 1 |
| Cluster_8 | chr8 | 124405254 | 124405307 | 1 |

St. Jude - Confidential

**Supplementary Table S3. Oligonucleotide sequences used in this study.**

List of all primers and sgRNA sequences used for genotyping, cloning, CRISPR-Cas9 editing, and sgRNA library screening.

| **Primer** | **Sequence 5'> 3'** |
|---|---|
| Ostir1 F | TGATAATATGGCCACATGACATACTTTCCTGAAGAGGTCG |
| Ostir1 R | TTAGTGGCTCCGCTTCCTGATCTCAGAATCTTCACA |
| P2A-GFP F | AAGCGGAGCCACTAACTTCTCCC |
| P2A-GFP R | TTGGCTGCAGGTCGATTATACCTTACGCTTCTTCTTTGGCTTGTAC |
| HA miniAID F | ACTCGAACAGCTTCGAAACTCTGGTGCATAC |
| HA miniAID R | GGGGGGGGCGGAATTTTATTTATACATCCTCAAATCGAT |
| CFP F | CGGTGCCTGAACGCGCTCGAGCGGGATCAATTCCG |
| CFP R | TGTCGACTTAACGCGTTACTTGTACAGCTCGTCCATG |
| sgMyc F | CACCGCTGTACGGAGTCGTAGTCG |
| sgMyc R | AAACCGACTACGACTCCGTACAGC |
| sgCcl2-1 F | CACCCACCTGGCTGAGCCAACACG |
| sgCcl2-1 R | AAACCGTGTTGGCTCAGCCAGGTG |
| sgCcl2-2 F | CACCGCAAGATGATCCCAATGAGT |
| sgCcl2-2 R | AAACACTCATTGGGATCATCTTGC |
| sgNf1-1 F | CACCGGAAACGTGGCATGTCTCGG |
| sgNf1-1 R | AAACCCGAGACATGCCACGTTTCC |
| sgNf1-4 F | CACCGGACGAGAGCAACATAAACA |
| sgNf1-4 R | AAACTGTTTATGTTGCTCTCGTCC |
| sgCxcl1-F | CACCGCAGTGGCGAGACCTACCTG |
| sgCxcl1-R | AAACCAGGTAGGTCTCGCCACTGC |
| sgRNA screen first round PCR primer F | TCGTCGGCAGCGTCAGATGTGTATAAGAGACAGAATGGACTAT CATATGCTTACCGTAACTTGAAAGTATTTCG |
| sgRNA screen first round PCR primer R | GTCTCGTGGGCTCGGAGATGTGTATAAGAGACAGCTTTAGTTT GTATGTCTGTTGCTATTATGTCTACTATTCTTTC |
| index primer N701 | TCGCCTTATAAGGCGA |
| index primer S504 | AGAGTAGAAGAGTAGA |
| index primer N704 | GCTCAGGATCCTGAGC |
| index primer S507 | AAGGAGTAAAGGAGTA |

| | |
|---|---|
| Cre-ERT2 genotyping primer F | TTAATCCATATTGGCAGAACGAAAACG |
| Cre-ERT2 genotyping primer R | CAGGCTAAGTGCCTTCTCTACA |
| Rosa26 WT F | TTCCCTCGTGATCTGCAACTC |
| Rosa26 WT R | CTTTAAGCCTGCCCAGAAGACT |
| LoxP-Stop-LoxP-tdTomato genotyping primer F | CGAAGTTATATTAAGGGTTCCGGATCA |
| LoxP-Stop-LoxP-tdTomato genotyping primer R | CCTTCGCTGCGGTCTTG |
| Trp53-null genotyping primer F | TGTTTTGCCAAGTTCTAATTCCATCAGA |
| Trp53-null genotyping primer R | TTGTAGTGGATGGTGGTATACTCAGA |
| Trp53 WT genotyping primer F | GTGAGGTAGGGAGCGACTTC |
| Trp53 WT genotyping primer R | TTGTAGTGGATGGTGGTATACTCAGA |

#### Supplementary Methods

##### *In vivo* bioluminescence imaging

OS cells were infected with lentiviruses co-expressing luciferase and the yellow fluorescent protein (YFP)(VCL20SF2-Luc2aYFP)(1) and sorted for the $YFP^+$ population. About 3 × $10^6$ cells were resuspended in 100 µL of PBS and injected subcutaneously into the right flank or intravenously into the tail vein of recipient mice. Flank tumor growth and metastasis to the lungs (with flank shielding; one image per side) were assessed at the indicated time points by bioluminescence imaging using an IVIS Spectrum or IVIS-200 system. Five to ten minutes prior to imaging, animals were injected intraperitoneally with D-Luciferin (15 mg/mL in sterile saline) at a dose of 150 mg/kg. After drug administration, animals were anesthetized using isoflurane and maintained via nosecone on a heated imaging bed within the system for the duration of the scan. Following imaging, animals were allowed to recover on a heating blanket under observation and supplemented with oxygen as required. All imaging signals were normalized at the end of the entire study and quantified for presentation and statistical analysis.

##### Histology and Immunohistochemistry

Tissues were fixed in 10% formalin, bone tumors were decalcified in 10% EDTA, embedded in paraffin, and sectioned at 5 µm. Serial sections were obtained. One slide was stained with Hematoxylin and Eosin (H&E) while the adjacent slides were used for immunohistochemistry using the following antibodies raised against MYC (#ab32072; Abcam), SATB2 (#384R-18; Cell Marque), CD3 (#sc-1127; Santa Cruz), F4/80 (#70076; Cell Signaling), CD11c (#97585; Cell signaling), B220 (#553084; BD Biosciences), and CD161c (#39197; Cell Signaling). Stained slides were scanned with a Panoramic 250 FLASH III Digital Scanner and analyzed using HALO Link and ImageJ.

##### Complete blood count (CBC)

Mouse peripheral blood was collected from the submandibular vein into a 1.5 mL tube containing 10% EDTA, then mixed to prevent clotting. The blood samples were analyzed by the St. Jude In-house Pathology Core Facility using automated methods.

##### Immunoblotting

Cells were lysed in RIPA buffer, and protein lysates were used for electrophoresis on an SDS-PAGE gel (#NP0335BOX; Thermo Fisher Scientific), followed by protein transfer to a PVDF membrane (#IPVH00010; MilliporeSigma) at 100 V for 1 hour. Membranes were blocked with 5% non-fat milk in TBS-T (10 mM Tris, pH 8.0, 150 mM NaCl, 0.5% Tween-20) for 1 hour at room temperature, then incubated overnight with primary antibodies at 4 °C with gentle rocking, including GAPDH (#AM4300; Thermo Fisher Scientific; 1:5,000), MYC (#C15410210-50; Diagenode; 1:2,000), HSC70 (#sc-7298; Santa Cruz; 1:500), HA (#3724S; Cell Signaling; 1:2,000), and Cas9 (#632607; Takara; 1:2,000). Membranes were washed three times in TBS-T and then incubated with corresponding secondary antibodies [donkey anti-rabbit IgG HRP (#NA934; GE Healthcare; 1:5,000) or sheep anti-mouse IgG HRP (#NA931; GE Healthcare; 1:5,000) in 5 % non-fat milk/TBS-T for 1 hour at room temperature. After three washes in TBS-T, blots were developed with ECL (#NEL1005001; PerkinElmer) and visualized by using the LOR System.

**Competitive proliferation assay (CPA)**

Lenti-Cas9-BSD was used to generate Cas9-expressing OS770 cell lines. Infected OS770 cells were selected by Blasticidin S (BSD, 10 μg/mL, #A1113903; Thermo Fisher Scientific). Cas9-expressing OS770 cells were transduced with Lenti-sgMyc-Puro-IRES-CFP or Lenti-sgNT-Puro-IRES-CFP. Half of the infected cells were selected by puromycin (1 μg/mL, #ant-pr-1; InvivoGen) and collected for immunoblotting or genomic DNA extraction to determine indel frequency by Sanger sequencing. The fitness of the other half was assessed by measuring the percentage of CFP-positive cells at days 3, 6, 9, and 13 post-infection by flow cytometry. Data analysis and presentation were produced by FlowJo software. The CFP-positive percentage was normalized to the starting time point and analyzed using GraphPad Prism.

**Spectral Karyotype (SKY)**

OS cells were harvested after a four-hour colcemid incubation to arrest actively dividing cells in metaphase. Spectral karyotyping was performed using the mouse SkyPaint DNA kit (#FPRPR0030; Applied Spectral Imaging) and a concentrated antibody detection (CAD) kit (#FPRPR0033; Applied Spectral Imaging) following the manufacturer's instructions. Images were acquired with a Nikon Eclipse E600 fluorescence microscope equipped with an interferometer (Spectra Cube: Applied Spectral Imaging) and a custom-designed filter cube (Chroma Technology Corporation, Rockingham, VT). SKY analysis was performed using HiSKY software version 8.4 (Applied Spectral Imaging). Chromosomes were labeled with different probes and displayed in distinct colors.

#### RNA sequencing

Established OS770 and OS329 cells were collected for total RNA extraction when the culture reached approximately 60% confluency. OS770 Cas9-expressing cells were infected with Lenti-puromycin-U6-sgRNA against the coding regions of *Ccl2*, and total RNAs were collected from puromycin-selected cells at day 5 when the culture reached approximately 60% confluency. Total RNA was extracted from replicate samples using TRIzol (#15596026; Thermo Fisher Scientific). About 200 ng of total RNA was treated using Kapa rRNA depletion reagents to remove ribosomal RNA, followed by conversion into cDNA libraries using Kapa RNA HyperPrep Kit with RiboErase (#KK8561; HMR).

#### Single-cell sequencing

All samples, including freshly enriched GNP fraction, cultured sphere cells at passage 8, and sphere cells overexpressing ectopic Myc, were dissociated into single-cell suspensions and sorted by flow cytometry for live singlets prior to processing with the 10x Genomics Chromium Next GEMM Single Cell 3' GEM v3.1 (#1000121; 10x Genomics) chemistry, according to the manufacturer's protocol, with a targeted recovery of around 7,000 cells. Microfluidics chips were run on the 10 × Genomics Chromium machine. Final libraries were run on an Illumina NovaSeq 6000 with the following parameters per sample: 300 million reads from the 100 bp paired-end sequencing pipeline.

#### Whole-genome sequencing

Genomic DNA was extracted from OS cells using a PureLink Genomic DNA extraction Kit (#K182002; Invitrogen) according to the manufacturer's protocol. The genomic DNA was submitted to St. Jude's in-house Hartwell Center for library preparation and sequencing using the Illumina NovaSeq platform. Sequencing reads were aligned to the mouse reference genome (mm10) using BWA. PCR duplicates were removed using Picard, and variant calling was performed using the Genome Analysis Toolkit (GATK)(2).

#### Cytokine array

About 0.5 × $10^6$ OS770 or OS329 cells were plated and cultured for two days. 1 mL of supernatant cultured medium was collected for the detection of secreted cytokines and chemokines using the Proteome Profiler Mouse Cytokine Array Kit, Panel A (#ARY006; R&D Systems), following the manufacturer's instructions.

**Cross-species transcriptome comparison**

RNA-seq expression data from OS770 and OS329 cells were compared with bulk RNA-seq profiles from 88 pediatric OS patients in the Therapeutically Applicable Research to Generate Effective Treatments (TARGET) OS cohort, obtained from the NCI Genomic Data Commons (GDC)(3). Mouse gene-level TPM expression values and human STAR-aligned TPM values were independently $\log_2$-transformed (x+1). To enable cross-species comparison, high-confidence one-to-one orthologous gene pairs were retrieved from Ensembl BioMart (release 113, GRCm39/GRCh38) using the mmusculus_gene_ensembl dataset, filtering for ortholog_one2one orthology type with a confidence score of 1. After intersecting with genes detected in both expression matrices, orthologous gene pairs (14,926 for OS770 and 13,587 for OS329) were retained. To focus on expressed genes, pairs with a mouse mean $\log_2(TPM+1) < 1$ were excluded. Mean expression per gene was computed across replicates (mouse) or across all patient samples (human), and Pearson and Spearman correlations were calculated between the two species. A linear regression line was fitted to the filtered gene set.

**OS cohorts Kaplan–Meier analysis**

For the *NCR1* gene expression Kaplan–Meier analysis, the R2 database (http://hgserver1.amc.nl) was used to generate the Kaplan–Meier survival curves for patients with OS using the Mixed Osteosarcoma (Mesenchymal) Kuijjer-127-vst-ilmnhwg6v2 dataset. Patients were stratified into high- and low-*NCR1* expression groups based on an automated scan cut-off (expression value = 168.7) for the log-rank test. For the NK signature gene expression Kaplan–Meier analysis, a curated set of 23 genes associated with NK cell identity and cytotoxic function was selected for survival analysis: *NCR1, NCR3, KLRB1, KLRD1, KLRK1, KLRC1, KLRC2, GNLY, NKG7, PRF1, GZMB, GZMK, GZMH, IFNG, NCAM1, FCGR3A, CD247, TYROBP, FCER1G, XCL1, XCL2, CXCR3*, and *CX3CR1*. Three independent OS cohorts were analyzed. RNA-sequencing data and matched clinical annotations for the TARGET-OS cohort (n = 86) were obtained from the NCI Genomic Data Commons (GDC) via the GDC REST API (3). Gene expression was quantified as $\log_2(TPM + 1)$ from STAR-Counts files (unstranded TPM column). Two microarray cohorts were retrieved from the NCBI Gene Expression Omnibus (GEO): GSE21257 (n = 53)(4) and GSE39055 (n = 37)(5). For microarray data, probe-to-gene mapping was performed using the platform annotation table; when multiple probes mapped to the same gene symbol, the probe with the highest expression value per sample was retained. CD247 was absent from the GSE21257 analysis (22/23 genes evaluated); XCL2 was absent from the GSE39055 analysis (22/23 genes evaluated). For each gene and cohort, patients were stratified into high- and low-expression groups using the cohort-specific median as the cutoff, yielding equal group sizes of 43/43 (TARGET-OS), 27/26 (GSE21257), and 19/18 (GSE39055). Survival distributions were estimated by the Kaplan–Meier method and compared between groups using the log-rank test.

**_CCL2_ and _CCR2_ expression analysis across OS patient cohorts**

Single-cell RNA-seq data from 11 human OS patients (eight osteoblastic, three chondroblastic; six primary tumors, two recurrent lesions, three lung metastases) were obtained from the GSE152048 as previously described (6). Briefly, raw reads were aligned to GRCh38 using Cell Ranger (v2.1.1); cells with fewer than 300 detected genes or mitochondrial content exceeding 10% were excluded, and doublets were removed using DoubletFinder (v2.0.2), yielding 100,987 high-quality transcriptomes. Cell type annotation was performed by the original authors based on canonical marker genes, identifying 11 major clusters including osteoblastic OS cells, T cells, NK cells, myeloid cells, fibroblasts, and endothelial cells, among others. For the present study, *CCL2* and *CCR2* expression was queried across annotated cell clusters to characterize the cellular sources of CCL2 and the distribution of its cognate receptor CCR2 within the OS tumor microenvironment. Gene expression data from OS patient biopsies were obtained from the GEO database (GSE21257)(4), which comprises 53 pre-treatment primary tumor biopsy samples profiled on the Illumina HumanWG-6 v3.0 Expression BeadChip (GPL10295). Samples were classified into two groups based on clinical outcome metadata: patients who developed metastasis at any time point (n = 34) and patients who remained metastasis-free (n = 19). Probe-level expression values were extracted for *CCL2* (probe: cSj1OhRehCQFQHWCH4) and *CCR2* (three probes: N103wwHkAJ4Ln4eE.E, NOTzc4znDM7M5DMA.I, Kn2SO1_OdM_9XD_Uuo), as annotated in the GPL10295 platform file. For *CCR2*, expression was summarized as the mean across all three probes per sample. All expression values are reported as $\log_2$-transformed fluorescence intensities as provided in the original dataset without additional normalization. Between-group comparisons were performed using the Welch two-sample t-test.
